# Exploring Quorum-Sensing-Based Bacterial GMOs for Addressing Bovine Mastitis: A Developmental Approach

**DOI:** 10.1101/2023.08.07.552207

**Authors:** Adrija Adhikary, Abeg Dutta, Udit Ghosh, Sourin Chatterjee, Shreyash Borkar, Shubhamay Das

**Affiliations:** Indian Institute of Science Education and Research, Department of Biological Sciences, Kolkata, India; Indian Institute of Science Education and Research, Department of Mathematics and Statistics, Kolkata, India

**Keywords:** Autoinducing Peptide I (AIP-I), Genetically Modified Organism (GMO), Bovine Mastitis (BM), Nisin, DNase I, Quorum sensing (QS), Biofilm

## Abstract

Bovine mastitis, a prevalent and costly disease affecting dairy cows, severely impacts milk production, quality, and cow health and poses significant challenges to the dairy industry worldwide. Traditional treatment strategies relying on antibiotics face concerns of antibiotic resistance and residual antibiotic residues in dairy products. Therefore, there is an urgent need for innovative and sustainable approaches. Our study undertakes a preliminary investigation into the formulation and efficacy of a GMO-based antibiotic-free treatment in fighting Bovine mastitis. We propose to use the bioengineered defensin Nisin PV as the antimicrobial to specifically target the causative pathogen *Staphylococcus aureus*. To disrupt the biofilm formation, we have leveraged the potency of DNaseI. By hacking the quorum sensing technique of *Staphylococcus aureus*, our proposed genetically modified bacteria would sense the presence of pathogens and initiate their eradication by producing Nisin PV and DNaseI. Results show the efficacy of DNase I against bacterial biofilm formation, in addition to the efficacy of our proposed sensor, which is based on the agr quorum sensing system of *Staphylococcus aureus*, a major cause of bovine mastitis. Molecular dynamics and simulations show that our choice of the bacterial defensin, Nisin PV, is less susceptible to cleavage by bacterial strains resistant to the native bacteriocin Nisin A. Additionally, we theoretically propose a lysis-based kill switch to ensure the release of the therapeutic components. The solution proposed in this study has the potential to offer a sustainable and effective alternative to antibiotic-based treatments. Implementation of bioengineered defensins could reduce disease incidence, minimize economic losses, and promote responsible antimicrobial stewardship in the dairy industry.

## 1 INTRODUCTION

Bovine mastitis, an inflammatory disease of the mammary glands, continues to pose significant challenges to the dairy industry worldwide. It is one of the most prevalent and costly diseases affecting dairy cows, leading to substantial economic losses and compromised milk quality (Halasa et al., 2007). As per a study conducted in the central region of India associated with dairy animals suffering from subclinical bovine mastitis, it was analyzed that 48.53% and 36.57% of the overall loss per lactation were due to the reduction in milk quantity due to disease progression and costs incurred by veterinary treatment, respectively. Based on population statistics and the incidence of mastitis seen in the study, the researchers projected the annual economic losses to be more than thousands of dollars for Maharashtra and Pune alone (Sinha et al., 2014). In another study constituting 86% of the dairy cow population of the United States, it was noted that more than 350 kg of milk loss was suffered by each cow in a year due to mastitis (Blosser, 1979). A comprehensive understanding of the economic implications of mastitis based on a model assessed on an average dairy farm in the Netherlands calculated the average losses as C17–198 per cow in a year (Huijps et al., 2008). In biological terms, mastitis is defined as inflammation of the parenchyma of the mammary glands. Initially, the pathogens infect the udder tissue, multiply, and finally attack the host immune system, leading to such inflammation (Thompson-Crispi et al., 2014). Often referred to as the silent killer of cows, bovine mastitis can occur in four stages, namely, infection, subclinical bovine mastitis, clinical bovine mastitis, and recurrent clinical mastitis. The progression of the disease to the clinical and further stages makes the treatment of bovines extremely difficult (Girma and Tamir, 2022). A better approach would be to treat the disease at an early stage. In this study, we frame a potential novel approach to harness an inherent property of the causative organism for treating the disease at the early subclinical stage.

Due to its lack of visible symptoms, subclinical bovine mastitis renders no apparent physical signs of inflammation or abnormality in the affected udder. Despite being conventionally undetectable, it has a major impact on the ability of the affected cows to produce milk, the quality of their milk, and their general health. The detrimental effects on milk production, milk quality, and human nutrition can be lessened with early detection and effective therapeutic strategies for subclinical mastitis (Paramasivam et al., 2022). Despite the presence of several causative pathogens, bacterial pathogens plays the largest role. *Staphylococcus aureus* and *Streptococcus agalactiae* are contagious pathogens, characterized by subclinical infections and adaptive to the udder environment. On the other hand, *Streptococcus uberis* and *Escherichia coli* are considered environmental pathogens (Kibebew, 2017). A study conducted in Ethiopia also confirmed that the majority of the isolates, i.e., 44.9%, were positive for *Staphylococcus aureus*. Other species, such as, *Streptococcus* spp. and gram-negative *Escherichia coli* followed the list with an abundance of 25.3% and 8.8% respectively (Birhanu et al., 2017)

However, traditional treatment strategies, relying heavily on antibiotics, have encountered growing concerns due to the emergence of antibiotic-resistant bacteria and the potential for residual antibiotic residues in dairy products. *Staphylococcus aureus* has also become increasingly antibiotic-resistant to different spectrums of antibiotics, which has further aggravated the situation (for Disease Control et al., 2002; Guo et al., 2020). The acquisition of resistance genes by horizontal gene transfer is one of the primary methods for antibiotic resistance in *Staphylococcus aureus*. In addition to it, *Staphylococcus aureus* can create biofilms, which are complex bacterial populations encased in a protective matrix (Lister and Horswill, 2014). Antibiotic penetration and efficacy can be hampered by biofilms, which act as a physical barrier. Antibiotic usage also puts *Staphylococcus aureus* populations under selection pressure. Antibiotic use over an extended period of time may encourage the survival and spread of resistant populations that have developed resistance mechanisms. This can happen by selectively eliminating the bacteria that are susceptible, allowing the resistant ones to flourish and proliferate (Águila-Arcos et al., 2017; Lowy et al., 2003). Consequently, there is an urgent need for innovative and sustainable approaches to combat bovine mastitis.

In recent years, bioengineering has emerged as a promising field for the development of novel therapeutic strategies (Field et al., 2015; Cruz et al., 2022). Defensins have garnered substantial attention among the various bioengineered molecules due to their potent antimicrobial properties and significant roles in innate immunity. Naturally occurring defensins, small cationic peptides, exhibit broad-spectrum antimicrobial activity against a wide range of pathogens, including bacteria, fungi, and viruses. Their ability to rapidly kill invading microorganisms while modulating the host immune response makes them ideal candidates for combating infectious diseases (Beńıtez-Chao et al., 2021).

In our approach, we propose to exploit the inherent quorum-sensing technique of *Staphylococcus aureus* for sensing the presence of pathogens. The introduction of the two-component agrC-agrA signal transduction system into our proposed chassis, *Lactococcus lactis* LMG 7930, would enable it to sense the auto-inducing peptide I(AIP-I) molecules produced by the pathogenic *Staphylococcus aureus*. This strain was previously used as a potential probiotic and has been shown to reduce inflammation due to mastitis when injected into the udder (Armas et al., 2017). Following sensing, the killing of pathogens can be achieved using a combination of DNase I and Nisin PV, a bioengineered form of the bacteriocin Nisin A that is resistant to Nisin Resistance Protein (NSR) produced by *Streptococcus uberis* (Pérez-Ibarreche et al., 2021). These genes are then incorporated along with the quorum sensing unit to build the proposed genetic circuit. Expression of these proteins is regulated by the P2 promoter which is activated in the presence of AIP-I by phosphorylated agrA. The incorporation of a Lysis E7-based kill switch ensures the rupturing of the chassis organism and allows the involved gene products to be released out of the cell. Lysis E7 gene would be placed under a phosphorylated agrA sensitive weak promoter P3, such that the rate of production of Lysis E7 is lesser compared to DNase I and Nisin PV, allowing their accumulation prior to release by lysis. The agrC-agrA component is designed to be oriented opposite to the P2 and P3 promoters to prevent leaky gene expression. The gene circuit can be seen in Figure 1. The proposed strategy in this research holds great potential for revolutionizing the management of bovine mastitis.

**Figure 1.**
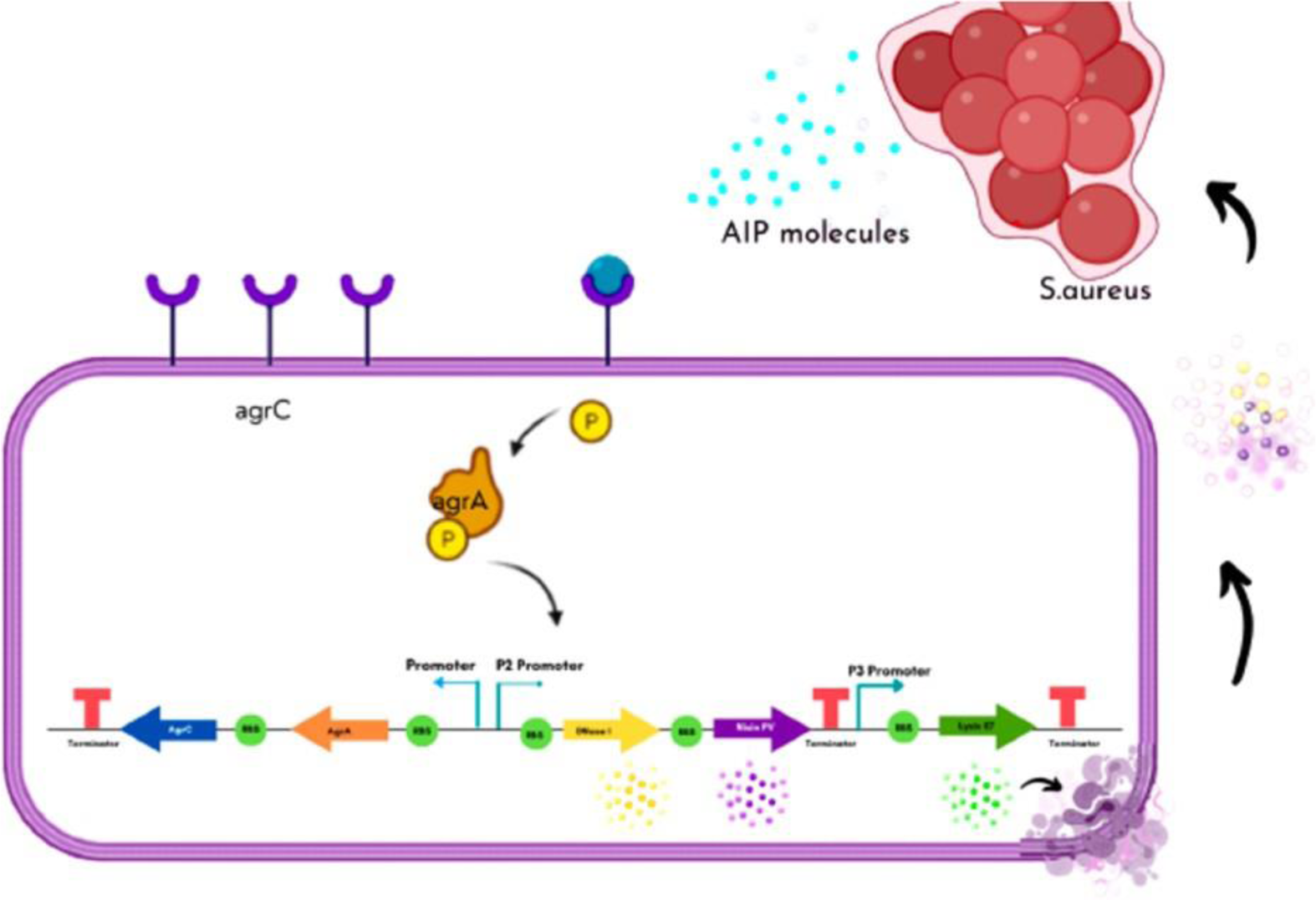
The working principle of the GMO. When the concentration of AIP-I exceeds a specific threshold as a result of the presence of Staphylococcus aureus cells, AIP-I molecules attach to agrC. Subsequently, the agrC-AIP-I complex functions as a kinase, leading to the phosphorylation of agrA. The phosphorylated agrA then binds to both P2 and P3, triggering the activation of Nisin PV and DNase I production downstream of P2, while Lysis E7 production occurs downstream of P3. However, the production rate of Nisin PV and DNase I surpasses that of Lysis E7 due to the heightened affinity of phosphorylated agrA for the P2 promoter compared to the P3 promoter. Therefore, our genetically modified organism will only undergo lysis after a substantial accumulation of Nisin PV and DNase I. If the AIP-I concentration is not above a certain threshold, then the GMO will not be switched on. RBS : Ribosome Binding Site.

By harnessing the power of bioengineering, we aim to develop a sustainable and effective alternative to conventional antibiotic-based treatment. The successful implementation of a bioengineered defensin as a therapeutic agent against bovine mastitis could significantly reduce disease incidence, minimize economic losses, and promote responsible antimicrobial stewardship in the dairy industry.

## 2 MATERIALS AND METHODS

### 2.1 Materials

All the resource material can be accessed from Supplementary Material.

### 2.2 Construction of the gene circuit

Cloning of all biological parts was done using a standardized cloning protocol with various steps elaborated as follows.

#### 2.2.1 Preparation of Competent Cells

A single colony was picked and inoculated in 10ml of Lysogeny Broth (LB). This was grown at 37°C overnight on a shaker. 1ml of this overnight culture was added to 100ml of prewarmed LB in a 500ml conical flask and grown on a shaker at 37°C till the OD reached 0.5. The culture was then cooled for 5 minutes on ice and transferred to clean falcon tubes. The cells were collected by centrifugation. The supernatant was discarded and the cells were placed on ice. TFB-I buffer (composition available in supplementary material) was used to resuspend the cells and they were inoculated on ice for 90 minutes. The cells were then collected by centrifugation (5 minutes,4000 g, 4°C). The supernatant was discarded and the cells were again incubated on ice. The cells were then resuspended in a TFB-2 buffer(composition available in supplementary material) (ice cold). 100*µ*l aliquots were prepared in 1.5ml centrifuge tubes and the cells were stored at -80°C.

#### 2.2.2 Plasmid Purification

Plasmids were extracted using Wizard® *Plus* SV Minipreps DNA Purification System Kit from Promega as per the manufacturer’s instructions.

#### 2.2.3 PCR Amplification

The PCR mix with the PCR tubes are placed on ice. Before starting the PCR program, all the PCR tubes are flicked and spun on a minispin centrifuge. The PCR program to be followed is as described in Figure 1 in the Supplementary material. The PCR mix used for amplification is described in Table 1.

**Table 1.**
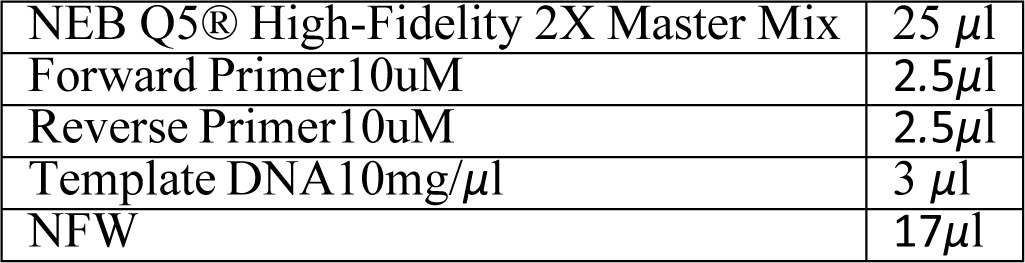
PCR mix for amplification.

|  |  |
| --- | --- |
| NEB Q5® High-Fidelity 2X Master Mix | 25 μl |
| Forward Primer10uM | 2.5μl |
| Reverse Primer10uM | 2.5μl |
| Template DNA10mg/μl | 3 μl |
| NFW | 17μl |

#### 2.2.4 PCR clean up

DNA purification from PCR mixture was done using Wizard SV Gel and PCR Clean-Up System Kit from Promega as per manufacturer’s instructions.

#### 2.2.5 Gel Extraction

DNA purification from agarose gel was done using Wizard SV Gel and PCR Clean-Up System Kit from Promega as per manufacturer’s instructions.

#### 2.2.6 Restriction digestion

The digestion mixture was prepared by combining the required components as per Table 2 and mixed gently by light flicking. After mixing, a short spin was performed to collect any liquid that may have collected on the sides. The digestion mixture was placed in a thermocycler and incubated for 5.5 hours at 37°C. Restriction enzymes from Promega were used.

**Table 2.** Mixture for restriction digestion.

|  |  |
| --- | --- |
| Restriction enzyme | 1μl each |
| Buffer(B) | 2 μl |
| BSA | 0.2μl |
| Template DNA | 1μg |
| MilliQ water | upto 20μl |

#### 2.2.7 Gel Electrophoresis

Agarose (1g) is mixed with 100ml of 1X TAE(composition available in supplementary material) to prepare a 1% Agarose gel. The mixture is then microwaved for 2-3 minutes until it completely dissolves, with occasional swirling. It is important to avoid over-boiling to prevent evaporation and spillage. The mixture is allowed to cool down to 50°C. EtBr is added to achieve a final concentration of 0.2-0.5 *µ*g/ml. The gel is cast in the gel tray and the comb is placed carefully. The gel is left at room temperature for 20-25 minutes until it solidifies properly. Loading dye is added to each sample. The gel box is filled with 1X TAE buffer, and the solidified agarose gel is placed in the gel box, ensuring it is fully covered with the buffer. If needed, an additional buffer is added. The ladder is loaded in the first lane, followed by careful loading of the samples. The gel is run at 80-120 Volts until the dye line reaches approximately 75-80% of the way down the gel (approximately 1-1.5 hours). The gel is then carefully removed and visualized using UV.

#### 2.2.8 Ligation

The components listed in the table 3 were assembled into a microcentrifuge tube placed on ice. It is crucial to add T4 DNA Ligase at the end. The molar ratio of insert to the plasmid, as determined using NEBioCalculator, is 3:1. The reaction is gently mixed by pipetting up and down, and then a brief microcentrifuge spin is performed. If the DNA fragments have cohesive (sticky) ends, the mixture is incubated at 16°C overnight. For blunt ends or single base overhangs as well, it is incubated at 16°C overnight. After the incubation, the mixture is heat-inactivated at 65°C for 10 minutes. The mixture is chilled on ice, and then 1-5 *µ*l of the reaction is transformed into 50 *µ*l of competent cells. The table 3 is for a 20ul Reaction.

**Table 3.**
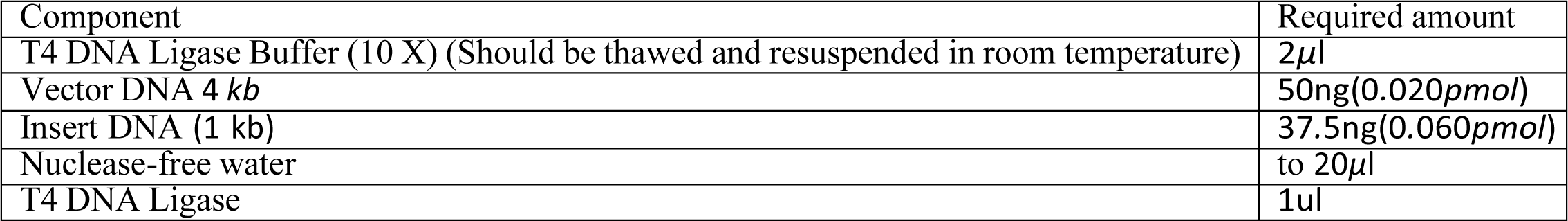
Mixture for ligation.

#### 2.2.9 Transformation

The competent cells (100*µ*l) are taken out from -80°C storage and placed on ice. The ligation mixture (10*µ*l) or miniprep Plasmids (1 or 2 *µ*l) are added to the 100*µ*l competent cells by pipetting, ensuring that the tip of the micropipette does not come in contact with the competent cells. The tube is closed and the contents are mixed by gently tapping on the side of the tube. The tube is kept on ice for 30 minutes. Meanwhile, the hot water bath is switched on and set to a temperature of 42°C, and the SOC media is placed in the incubator at 37°C. The tube is placed in a floating rack and heat shocked for 45 seconds at 42°C, using a timer for accurate time measurement. The tube is immediately transferred to ice and kept for 5 minutes. Afterward, SOC media is added to bring the volume up to 1000*µ*l, and the mixture is incubated in shaking conditions at 37°C for 1.5 hours. Meanwhile, the agar plates are kept at room temperature. After incubation, 100*µ*l is taken from the tube and spread with a cell spreader. The remaining contents of the tube are centrifuged at 10,000 x g for 3 minutes, the maximum supernatant is poured out, and the cell pellets are mixed with the leftover supernatant using a pipette and spread on another agar plate. The plate is then stored in an incubator at 37°C for 12-16 hours.

#### 2.2.10 Colony PCR

The PCR tubes were filled with 10*µ*l of Nuclease-Free Water (NFW) in each tube, as required. The culture plates with colonies were appropriately marked. A sterile tip was used to pick one colony and streak it at the designated position on a new plate. Subsequently, the same tip was placed in the PCR tube containing 10*µ*l of NFW. An equal number of autoclave PCR tubes were taken as the number of picked colonies and labeled accordingly. The reagents that were added to the PCR tubes to create a reaction mixture of 25*µ*l are in Table 4. The PCR tubes were spun on a minispin centrifuge before initiating the PCR reaction. The PCR program that was followed is as per Figure 1 available in the supplementary material.

**Table 4.** Mixture for colony PCR.

|  |  |
| --- | --- |
| Go Taq Green DNA master mix | 12.5μl |
| Template DNA | 5μl |
| VF2 | 1μl |
| VR | 1μl |
| NFW | 5.5μl |

### 2.3 DNase I biofilm assay

#### 2.3.1 Protocol for studying biofilm formation

The microtiter plate assay was used to observe bacterial adherence to a surface. Bacteria were incubated in 96-well plates, and after rinsing away the planktonic bacteria, the remaining adherent bacteria were stained with crystal violet dye. Henceforth, visualization or quantification of the stained biofilms was possible. To prepare the cultures, bacteria were inoculated into growth medium and grown overnight at 37°C. Pseudomonas cultures were diluted to 0.1 OD and added to the wells of a 96-well dish. After a 72-hour incubation period at 37°C, the cells were discarded, and the plate was washed with PBS to remove unattached cells and media components. 0.1% Crystal violet dye was then added, followed by incubation for 15 minutes and subsequent rinsing with PBS. The plate was dried, and the stained biofilms were solubilized using 33% acetic acid and incubated for 15 minutes. The solubilized dye was transferred to a new microtiter dish for measurement of absorbance at 595 nm using a plate reader, with 33% acetic acid as the blank.

#### 2.3.2 Standardisation of Biofilm assay

A standard Biofilm test in a 96-well plate was used to characterize DNase I. We chose to utilize *Pseudomonas aeruginosa* because of it being gram-negative and easily available. Before beginning the DNase I treatment, we had to standardize the technique by changing the experimental conditions to find the parameters that would produce the most biofilm. In our trials, we altered the media type, dilution, and incubation days. Following a review of the literature, we identified Luria broth (LB), Nutrient broth (NB), and Tryptone Soytone Broth (TSB), being suitable for the formation of biofilms. To find the optimal incubation period for biofilm growth in 96 well plates at 37°C, we first cultivated bacterial biofilm without DNase I in various media for various duration of incubation (24 hours and 72 hours). We also looked at how the yield of biofilm varied for two distinct secondary cultures prepared in 96-well plates. Before adding the primary culture to the 96-well plate, the optical density was diluted to between 0.1 and 0.05 OD with fresh medium. This was then incubated for the optimized 72 hours, according to the results of the previous experiment for various duration of incubation. Here, a variety of media were utilized (LB, NB, and TSB) to see whether the choice of media and dilution had any combined effects on biofilm yield. Biofilm quantification was carried out by measuring the absorbance at 595 nm. In conclusion, we standardized the 96-well plate biofilm assay by selecting NB media for bacterial growth, 0.1 OD, and 72-hour incubation for maximum biofilm output.

#### 2.3.3 DNase I-biofilm assay

To optimize the concentration of DNase I required for treatment, the effect of a gradient of DNase I in the range of 0–50 *µ*g/mL was studied. As a negative control for the experiment, uninoculated wells containing media and diluent were treated similarly and the readings obtained were subtracted from the test readings. Positive control wells containing inoculated media without DNase I were considered as “Control A595 nm”. The biofilm percentage reduction was then calculated as:

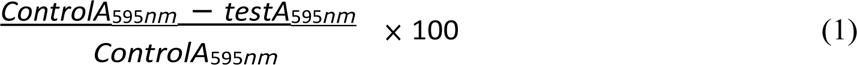

In order to determine the optimum concentration and time for DNase I treatment, we did an assay at two levels:

1. Pretreatment: It was done to determine the optimum concentration of DNase I which gives maximum degradation. A serial dilution of DNase I (0-25 *µ*g/ml) was prepared in NB media. Primary cultures were directly diluted to this media and added to 96 well plates as per the procedure mentioned in the above protocol. The layout used can be seen in Figure 2 available in supplementary material. Biofilm was allowed to develop in the presence of DNase1 for 72 hours. Plates were washed and processed as per the procedure mentioned in the above protocol. Absorbance was measured at 595 nm to quantify biofilm. Biofilm percentage reduction at different concentrations was compared and analyzed.
2. Posttreatment: This was done to determine the optimum time required for maximum biofilm degradation for the optimal concentration determined by pretreatment. Pseudomonas biofilm was allowed to develop in 96 well plates for 72 hours as per the protocol mentioned before. Well constituents, particularly bacteria, were then washed off in dH20. Media containing optimum concentration determined by pretreatment was then added into each well and incubated for different times ranging from 0-50 minutes. Plates were further processed by adding Crystal violet followed by Acetic acid. Biofilm quantification was carried out at 595nm. Biofilm percentage reduction was calculated and further analyzed.
3. Cloning and expression of DNase I: Initially, the gene fragment of the DNase I coding sequence with biobrick prefix and suffix which was synthetically obtained from Twistbioscience was amplified using Routine PCR and biobrick-specific primers. This was then ligated into the linearized plasmid backbone of PSB1C3 after restriction digestion by EcoRI and PstI following normal bio brick assembly. *E. Coli* DH5*α* was transformed and further verified using colony PCR using sequencing primers. The size of gene fragment after amplifying with sequencing primer was simulated as 1161 bp using Snapgene in-silico PCR simulator. For expressing the protein in *E. Coli* BL21, we cloned a composite part of DNase I, which consists of IPTG inducible Plac promoter(R0010) along with strong Ribosome Binding Site(B0034) and double terminator(B0015). This part was obtained synthetically from Twistbioscience. The gene fragment was amplified by Routine PCR using biobrick-specific primers and then ligated to the linearized plasmid backbone of PSB1C3 following normal biobrick assembly. Double digestion by EcoRI and PstI was done using optimized protocols for double digestion.*E. Coli* DH5*α* was initially transformed and transformants were screened and verified using colony PCR using sequencing primers. Snapgene in silico PCR simulator was used to determine the size of gene fragments after colony PCR. Further, the plasmids were extracted from transformed colonies and transformed *E. Coli* BL21 for expressing the protein. We did not get enough time to achieve protein purification due to COVID-19 restrictions and hence used industrially prepared bovine pancreatic DNase I.

**Figure 2.**
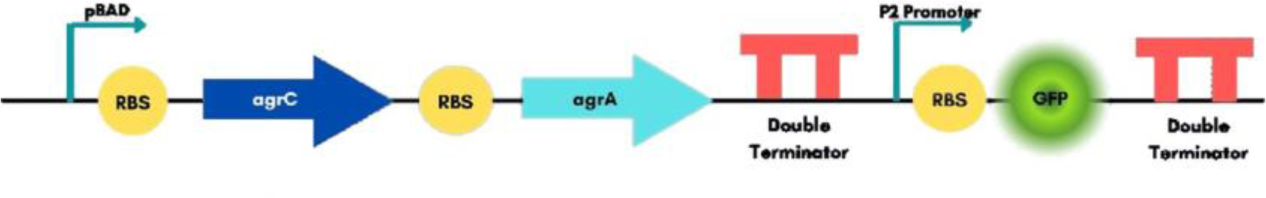
The regulatory circuit comprises of agrC and agrA genes, which are constitutively expressed via the pBAD promoter featuring a strong ribosomal binding site and a double terminator. P2 promoter downstream is activated in the presence of AgrA. Later to test if the AIP-I sensing module was working properly or not, a GFP reporter gene was incorporated downstream of the P2 promotor. The reasoning was that in the presence of AIP-I molecules, the P2 promoter could sense and activate the GFP gene which would then give an effective readout, hence confirming that the model senses AIP-I in its presence. RBS : Ribosome Binding Site.

**Figure 3.**
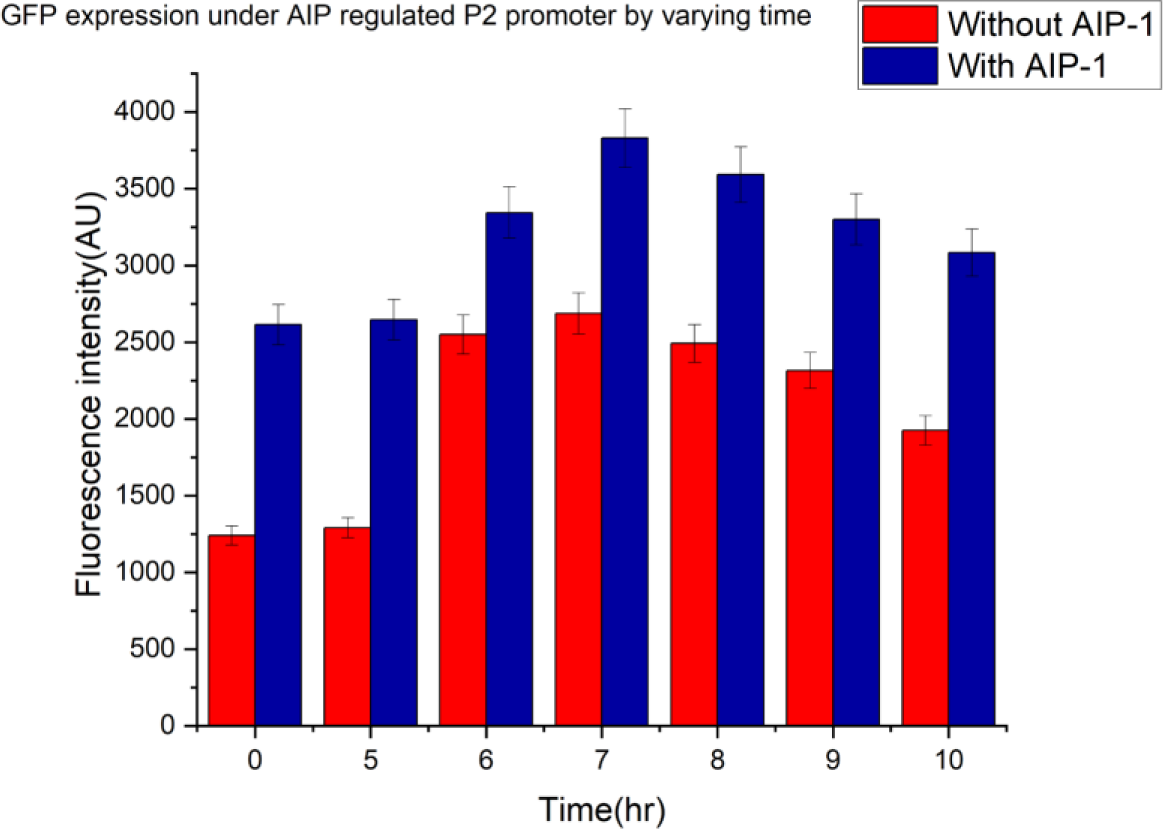
GFP expression with AIP-I alongside without AIP-I across varying time. The error bar represents the standard deviation of the results.

**Figure 4.**
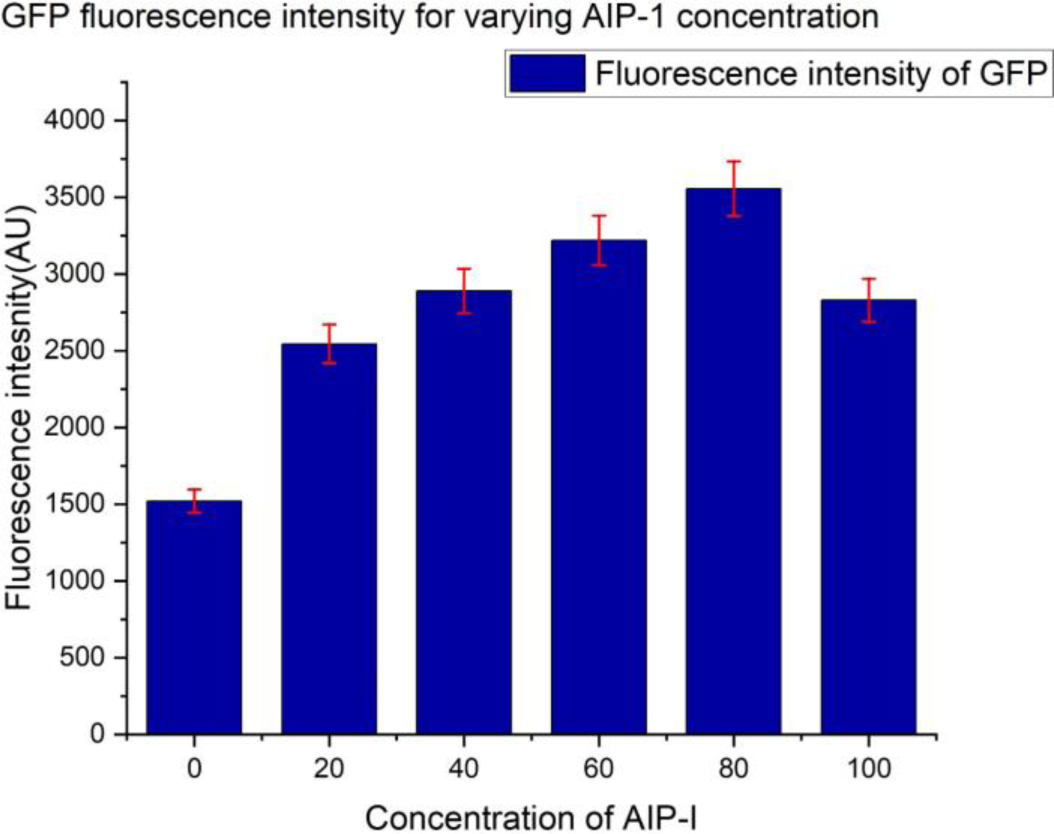
GFP expression with varying concentration of AIP-I. The error bar represents the standard deviation of the results.

### 2.4 Fluorescence assay to assess the improved AIP-I sensor

The assembly of 4 fragments (including the plasmid backbone) was performed using the NEB HiFi DNA Assembly kit. An LB agar plate containing bacteria with a unidirectional circuit and another LB agar plate with a bidirectional circuit containing bacteria were obtained. Separate primary cultures of both bacteria were provided in 10ml of LB agar supplemented with Chloramphenicol. The cultures were incubated overnight at 37°C. Autoclaved conical flasks (250 ml) were filled with 25 ml of fresh LB media (Containing chloramphenicol). Half of the flasks were inoculated with 1ml of the primary culture of unidirectional circuit bacteria, while the other half were inoculated with 1ml of the primary culture of bidirectional circuit bacteria. The flasks were incubated at 37°C until reaching an optical density (OD) of 0.6 at 600nm. Different concentrations of AIP-I molecules were added to induce the flasks. To assess fluorescence, a Polystyrene Nunc black 96-well fluorescence plate was acquired. Triplicates of 100*µ*l samples from each flask were added to the 96-well plate. LB media (100*µ*l) was added to each well for sample dilution. As a negative control, 200*µ*l of LB media was added to three wells. Fluorescence measurements were taken using a plate reader with an excitation wavelength of 395nm and a fluorescence wavelength measured of 509nm. The flasks were incubated at 30°C to allow for protein expression, and fluorescence was measured at various time points. Fluorescence from negative control(LB medium) was subtracted from the readings to obtain net fluorescence.

### 2.5 Molecular modeling

#### 2.5.1 Molecular docking and MM-PBSA/GBSA based free energy calculations

In order to characterize the efficacy of engineered nisin (Nisin PV) we first performed molecular docking of Nisin A and Nisin PV respectively with the Nisin Resistance Protein (NSR). Nisin A structure was mutated using PyMol and a PRO-VAL mutation was added to the 29th and 30th residues respectively to produce Nisin PV. These studies were inspired by the bioengineering nisin study(Field et al., 2019). Residues 22 to 34 from the N-terminus of the PDB structure with ID 1WCO were used as a representative of the segment of Nisin that enters the tunnel region of NSR for cleavage.

For the NSR (Nisin Resistance protein) molecule, the PDB ID 4Y68 was used. Waters associated with proteins were removed and polar hydrogens were added during pre-processing. Kollman charges were added to the protein. To determine a starting configuration for the NSR-nisin complex a docking program Autodock for ligand and protein binding was used where the Nisin (A/PV) was set as ligand and NSR was set as the receptor. We first performed the docking of Nisin A with the tunnel region of NSR and selected the lowest energy stage as the starting configuration. The docking location was found to be consistent with the ligand-binding site of Nisin as discussed in the literature (Field et al., 2019). The ligand-binding site for Nisin A was noted and Nisin PV was docked using the same coordinates.

The ensemble of poses generated from protein-protein molecular docking was fed into a Molecular mechanics Poisson–Boltzmann surface area (MM/PBSA) and molecular mechanics generalized Born surface area (MM/GBSA) implementation to perform binding free energy estimations and rescore the binding poses. We also performed residue-wise calculations for each Nisin subtype. We used the HawkDock server (Wang et al., 2019) for this purpose.

## 3 RESULTS

### 3.1 Detecting the presence of *Staphylococcus aureus* through AIP-I - sensor

In *Staphylococcus aureus*, which is the causative pathogen of Bovine Mastitis, agrA, and agrC are integral components of the agr quorum sensing system, which allows the bacteria to sense and respond to changes in population density(Thoendel and Horswill, 2013). agrC functions as the sensor histidine kinase, detecting and binding to specific autoinducing peptides produced by neighboring bacteria. Upon AIP-I binding, agrC undergoes autophosphorylation, activating agrA, the response regulator(Ji et al., 1995). agrA then acts as a transcription factor, modulating the expression of various genes involved in the quorum-sensing pathway. This includes the upregulation or downregulation of virulence factors, as well as the modulation of regulatory molecules like RNAIII and Rot. Through this regulatory network, agrA and agrC help coordinate the expression of virulence factors and other molecules crucial for the pathogenicity of *Staphylococcus aureus*, allowing the bacteria to respond to changes in population density and environmental conditions (Boisset et al., 2007).

Our strategy puts forward the idea of harnessing the intrinsic quorum-sensing mechanism of *Staphylococcus aureus* to detect the presence of pathogens. To achieve this, we propose to introduce the agrC-agrA two-component signal transduction system into our prospective engineered host, *Lactococcus lactis* LMG 7930. By incorporating this system, our chassis would gain the ability to sense and respond to auto-inducing peptide I(AIP-I) molecules produced by the pathogenic *Staphylococcus aureus*. This approach is a preliminary investigation into the development of Quorum-sensing based GMOs for targeted treatment of Bovine Mastitis and holds promise for the development of a reliable and efficient pathogen-sensing platform.

We designed a genetic circuit so that our GMO can sense the presence of AIP-I molecules in its surroundings and then regulate the downstream signaling. We designed a regulatory module that consists of agrC and agrA under a constitutive expression using pBAD promoter with a strong ribosomal binding site [iGEM registry: B0034] and a double terminator [iGEM registry: B0015]. This ensures the production of agrA and agrC protein and the detection of AIP-I molecules whenever present. In the presence of AIP-I, activated agrA binds to P2 promoter and increases the expression of downstream genes. Thus, we added a sensing module downstream of the regulatory module consisting of promoter P2. To test this sensing module, we incorporated *gfp* as a reporter gene downstream of the P2 promoter such that in the presence of AIP-I molecules P2 promoter can express GFP which can be observed in Figure2. So, we externally added AIP-I molecules to observe if the module is sensing its presence or not. To evaluate the functionality of our system, we conducted an exogenous supplementation of AIP-I molecules to ascertain the responsiveness of the engineered module. By externally introducing AIP-I, we aimed to investigate the system’s capability to detect and activate downstream genes to the AIP-I.

#### 3.1.1 AIP-I production

To activate the sensing module, we need AIP-I molecules, which will activate the agrC-agrA system thus increasing the promoter activity of P2. In *Staphylococcus aureus*, the genes agrB and agrD are responsible for the production of AIP-I molecules. Thus we cloned these two genes (AIP-I produce, iGEM registry: I746001) under Isopropyl ß-D-1-thiogalactopyranoside (IPTG) inducible pLac promoter(iGEM registry: R0010) in *E. coli* BL21. Protein expression was induced with IPTG. Protein expression was allowed for 10 hours at 30°C. The bacterial cells were pellet down and the supernatant containing the AIP-I molecule was filter-sterilized. The supernatant was lyophilized to increase the concentration of the content.

#### 3.1.2 Verifying the presence and activity of AIP-I

The presence of AIP-I in the transformed cells was confirmed through the assessment of GFP intensity generated by the P2 promoter within the AIP-Inducible GFP reporter gene circuit. The activation of the P2 promoter is initially mediated by agrA, which requires phosphorylation by agrC. agrC, in turn, becomes activated in the presence of AIP-I. To verify the presence of the protein, an *E. coli* BL21 expression vector containing a plasmid with the AIP-I receiver (agrA) and agrC system downstream of the pBAD promoter, along with GFP under the AIP-Inducible P2 promoter, was obtained. These bacterial cells were cultivated in two separate conical flasks containing LB media supplemented with Chloramphenicol. The cultures were incubated at 37°C until the optical density (OD) reached 0.6 at a wavelength of 600 nm. The AIP-I molecules were reconstituted by adding nuclease-free water. AIP-I was introduced into one flask, while the other flask remained without AIP-I. The expression of GFP was quantified by measuring the fluorescence using a fluorometer at the initial time point. Subsequently, the samples were incubated at 30°C for an additional 5 hours, with fluorescence measurements taken hourly from 5 to 10 hours. Figure3 clearly demonstrates a significant disparity in fluorescence intensity between the AIP-Inducible sample and the sample without AIP-I. This indicates successful expression of the AIP-I protein in our genetic circuit and confirms its activity.

#### 3.1.3 Assessing efficacy of AIP-I-sensor by Concentration-Dependent Activity

Furthermore, we investigated the concentration-dependent activity of the system by evaluating the expression of the GFP reporter construct in response to varying concentrations of AIP-I as shown in Figure4. For this experiment, *E. coli* BL21 cells expressing the AIP-I sensor with the reporter construct were incubated with AIP-I molecules reconstituted using nuclease-free water. Different volumes of the reconstituted AIP-I molecules were added to separate conical flasks to induce protein expression. The flasks were then incubated at 30°C for 5 hours. The GFP expression was quantified by measuring the fluorescence using a fluorometer. The obtained data were analyzed to establish the relationship between AIP-I concentration and GFP expression. The resulting analysis of the fluorescence data revealed that GFP expression increased with higher volumes of reconstituted AIP-I molecules. In other words, the concentration of AIP-I positively correlated with GFP expression in the media. This further substantiates the proper functioning of our sensing and regulatory module(agrC, agrA, P2, and GFP). Thus using our genetic module, we can express any set of genes in the presence of AIP-I.

It is important to note that at very high concentrations of AIP-I, the fluorescence intensity decreased, suggesting the occurrence of toxicity resulting from the elevated concentration of AIP-I.

### 3.2 DNaseI deters biofilm formation

After proving the efficacy of our sensing module(agrC-agrA-P2 system), we proceed to prove the potency of DNaseI, which is placed downstream of our P2 promoter. To identify the dosage concentration and optimal time for biofilm degradation, the activity of Bovine pancreatic DNase I (Sigma Aldrich) on biofilm production was assayed. *Pseudomonas aeruginosa* was used as a model biofilm-producing organism owing to its similarities to biofilm-forming pathogens, availability and it being a BSL-2 organism.

#### 3.2.1 Concentration dependant DNaseI assay

Biofilm assay was performed by varying DNaseI concentration and incubation time to identify optimal DNaseI dosage and time for biofilm degradation. To identify growth conditions to obtain maximal biofilm formation *Pseudomonas aeruginosa* was cultivated in different media, with varying starting inoculation and incubated for different duration at 37C. From Figure 5 biofilm formation in Nutrient broth(NB) media with incubation for 72 hours was found to be highest among the screened conditions and the biofilm yield with starting OD of 0.1 was observed to be always higher than 0.05 starting OD as seen in Figure 6.

**Figure 5.**
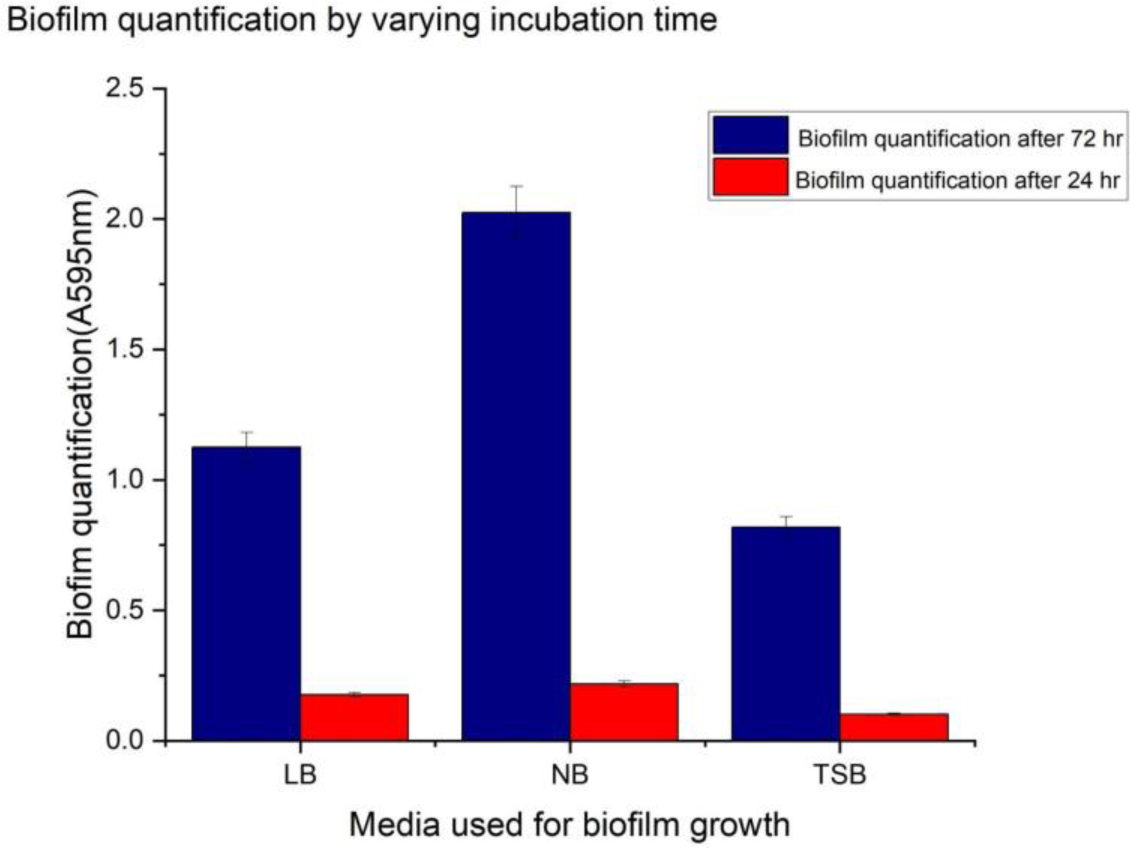
Biofilm yield in 96-well plate comparison in different growth media broths with incubation at 37°C for 72 and 24 hours. The error bar represents the standard deviation of the results.

**Figure 6.**
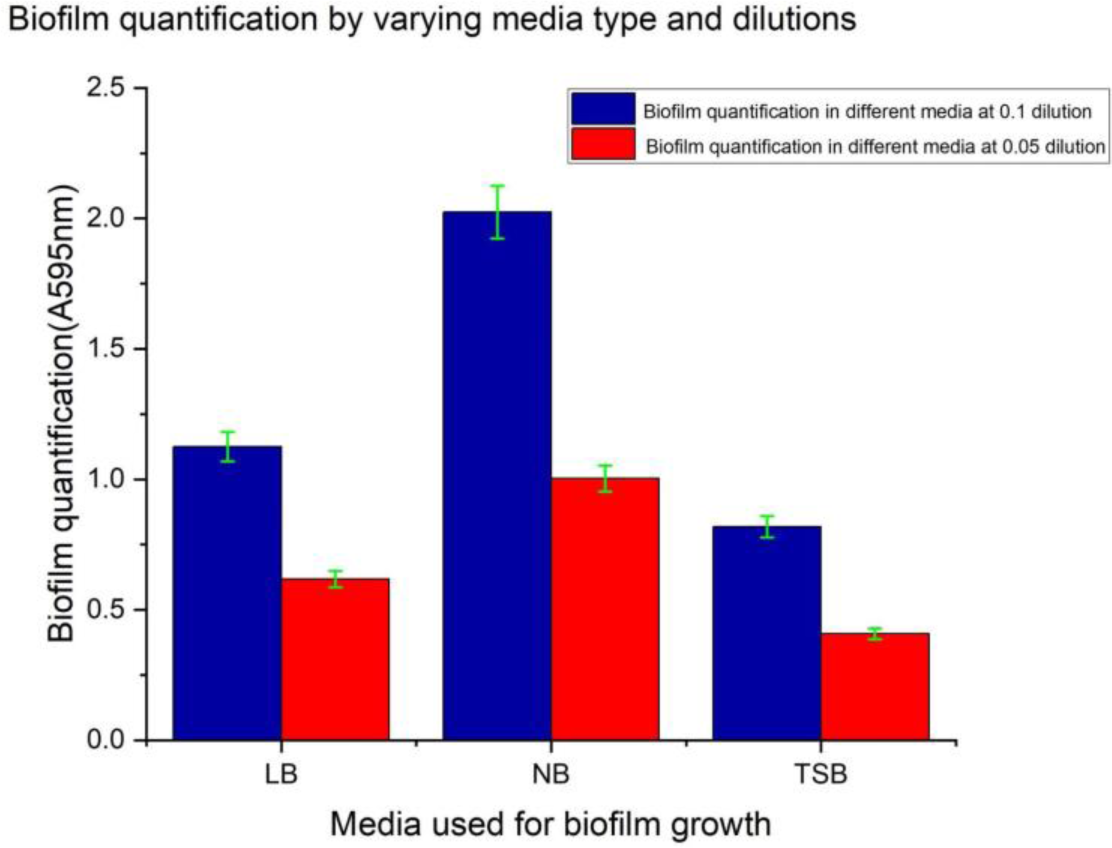
Biofilm yield comparison in different growth media with starting OD of 0.05 and 0.1 on incubation at 37°C for 72 hours. The error bar represents the standard deviation of the results.

The activity of DNaseI on biofilm was quantified in terms of biofilm percentage reduction (BPR) calculated by the following equation1.

Initially, the effect of DNaseI on biofilm formation was assayed by supplying different concentrations of DNaseI (0, 5, 10, 15, 20, 25 *µ*g/ml) in NB as seen in Figure 7. The biofilm was quantified after incubation at 37°C for 72 hours. 10 *µ*g/ml was able to reduce biofilm formation by 43%; subsequent increase in DNaseI concentration did not reduce biofilm formation. Subsequently, the ability of 10µg/ml DNaseI to degrade pre-formed biofilm was tested with varying treatment duration. The biofilm was allowed to form in NB for 72 hours at 37°C. The preformed biofilm was treated with 10 µg/ml DNaseI and incubated at 37°C for a different duration before quantifying the residual biofilm. Incubation for 40 minutes degraded 82% of bio-film. These can be seen in Figure 8. Hence, this experiment proves that DNaseI can degrade biofilm and supports the working hypothesis of DNaseI’s ability to degrade biofilm produced by *S. aureus*.

**Figure 7.**
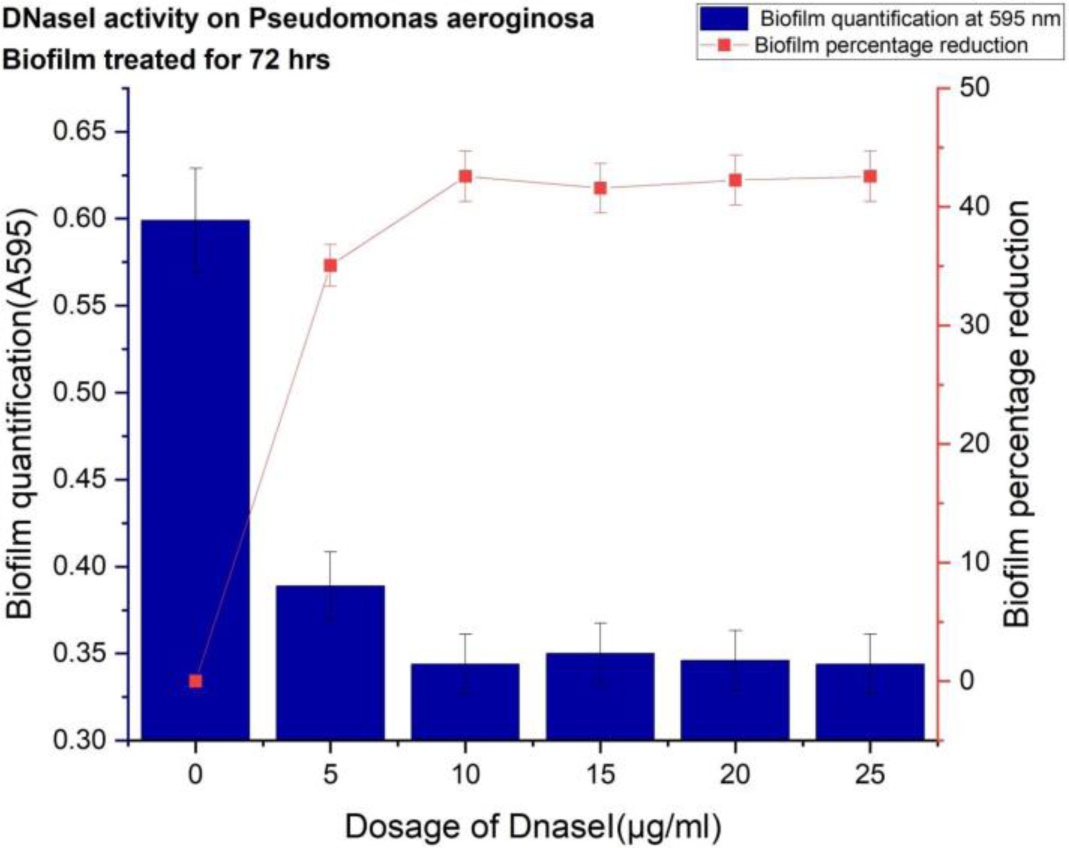
Biofilm formation in NB in the presence of different DNaseI concentrations after 72 hours incubation at 37°C.The error bar represents the standard deviation of the results.

**Figure 8.**
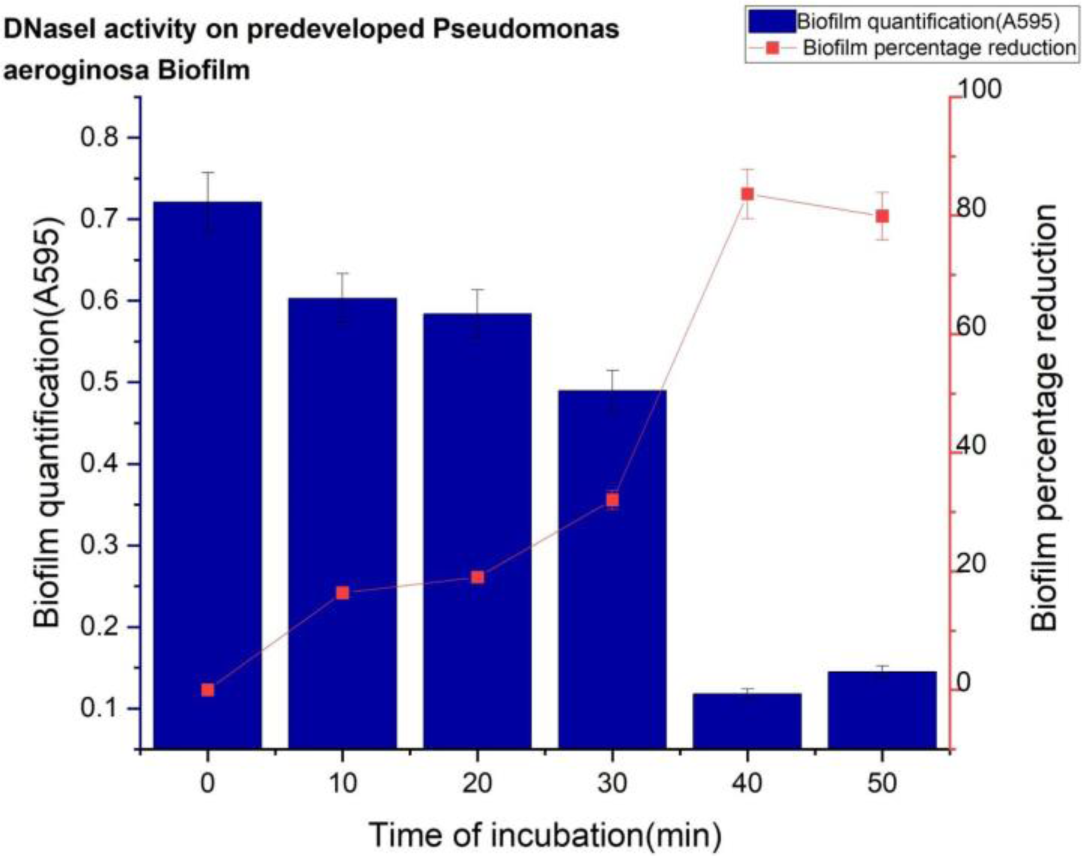
Quantification of biofilm degradation with time by treating pre-formed biofilm with 10µg/ml DNaseI and incubating at 37°C for different durations. The error bar represents the standard deviation of the results.

### 3.3 Molecular modeling

#### 3.3.1 Molecular Docking of Nisin with NSR

The docking results are summarized in Table 5. The highest magnitude pose corresponds to the binding of NisinA/PV to the tunnel region of NSR as shown in the figure 9.

**Table 5.**
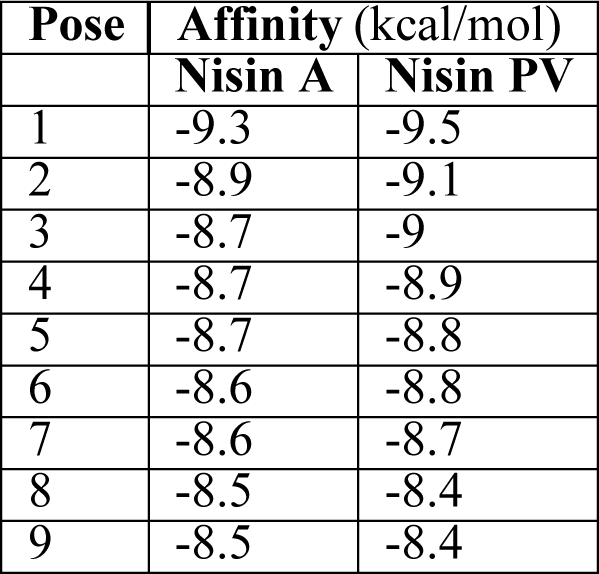
Affinity(Autodock score) of Nisin A vs Nisin PV for top 9 docking poses. The lowest free energy pose corresponds to the docking of Nisin in the tunnel region of NSR.

| Pose | Affinity (kcal/mol) |  |
| --- | --- | --- |
|  | Nisin A | Nisin PV |
| 1 | -9.3 | -9.5 |
| 2 | -8.9 | -9.1 |
| 3 | -8.7 | -9 |
| 4 | -8.7 | -8.9 |
| 5 | -8.7 | -8.8 |
| 6 | -8.6 | -8.8 |
| 7 | -8.6 | -8.7 |
| 8 | -8.5 | -8.4 |
| 9 | -8.5 | -8.4 |

**Figure 9.**
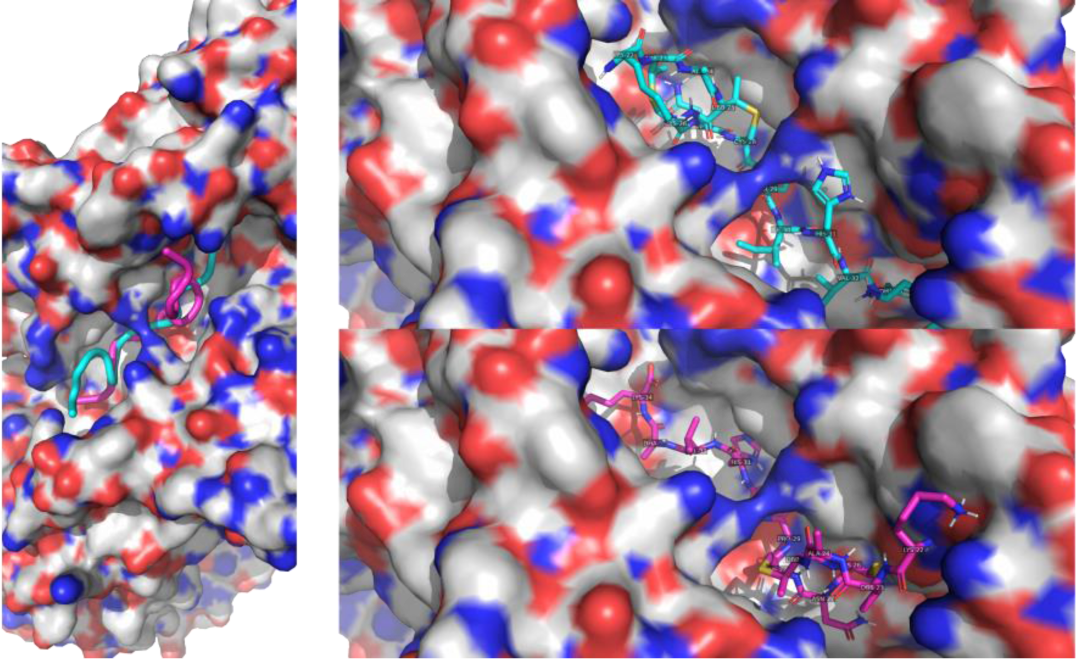
Docking of residues 22-34 N-terminus end of NisinA (cyan) and Nisin PV (magenta) to the tunnel region of NSR. Only the lowest free energy poses are shown.

**Figure 10.**
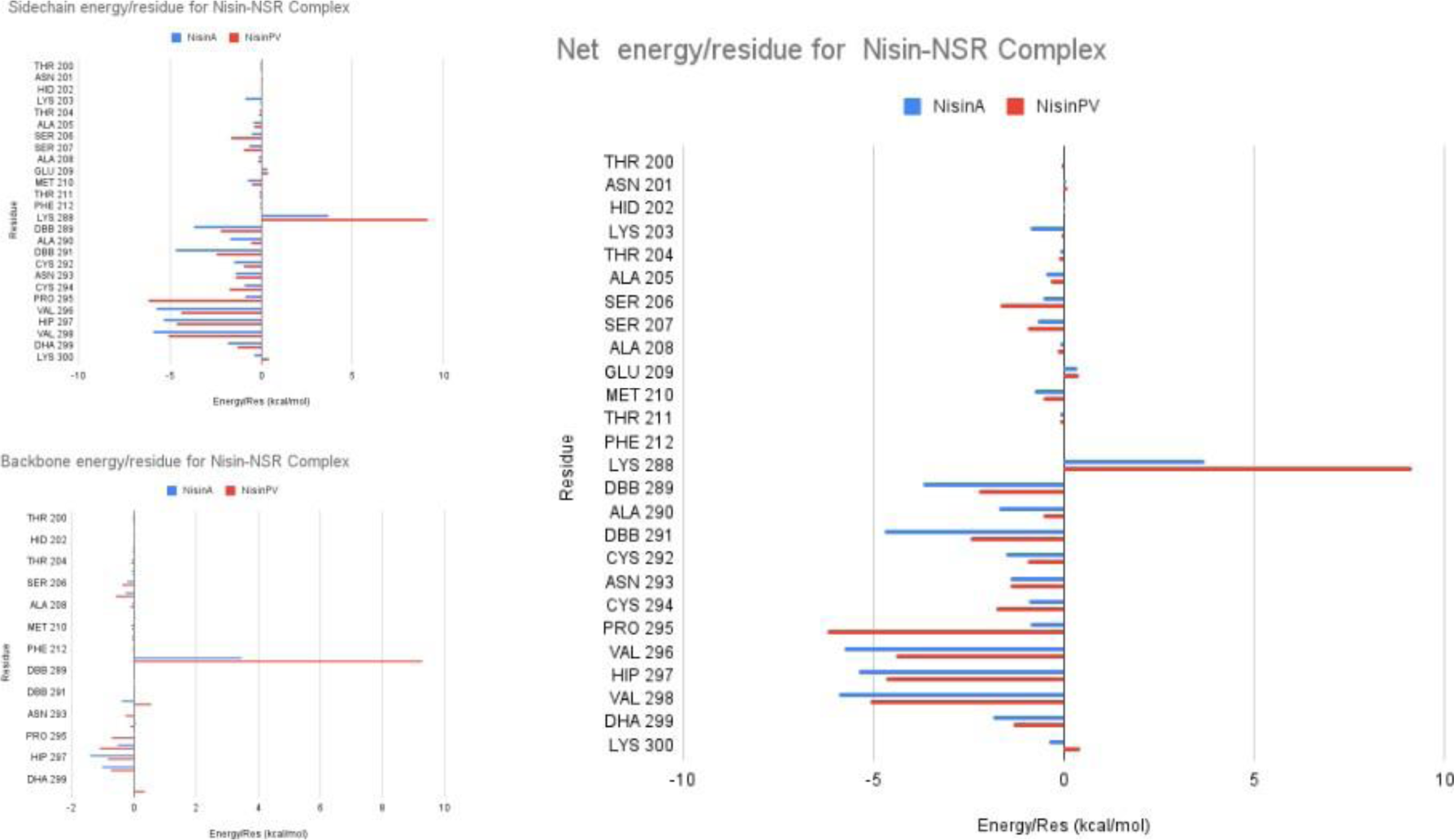
Residue-wise free energy for Nisin-NSR interaction.

#### 3.3.2 MM-PB/GB-SA based free energy calculations

MMPBSA calculations (10) reveal that at residue 29, the residue at which NSR is supposed to cleave Nisin, NisinPV shows significantly higher total free energy than the native Nisin indicating that cleavage of NisinPV is potentially more energetically expensive than cleavage of NisinA,

## 4 DISCUSSION

We have proposed a GMO-based therapy that targets the quorum-sensing system of *Staphylococcus aureus*, a common cause of bovine mastitis. In our approach, we target *Staphylococcus aureus* by exploiting its quorum sensing route and induce DNase I production in response to bacterial presence. Our novel strategy ensures that the GMO is only turned on when *Staphylococcus aureus* cells are present and the AIP-I concentration is greater than a certain threshold. Future improvements on our study can be undertaken to measure the exact AIP-1 concentration and standardize to check when the GMO works. The choice of promoter helps us to ensure that the GMO will not lyse until there has been a considerable buildup of DNase I and Nisin PV, which can break down the biofilm and make the bacteria more vulnerable to antimicrobial treatments. Due to the larger affinity of phosphorylated agrA towards the P2 promoter than the P3 promoter, the production rate of Nisin PV and DNase I is higher than the production rate of Lysis E7.

Instead of broad-spectrum antibiotics, this bacteriocin (Nisin PV) is more pathogen-specific. Our genetic circuit uses Lysis E7 to lyse our GMO for the effective release of the Nisin PV to target the pathogen *S. aureus*. This helps in the delivery of the bacteriocin as well as the killing of the GMO. Its crucial to note that Nisin is a crucial component of the quorum sensing system in *Lactococcus lactis*, where the presence of Nisin-A in the extracellular space triggers the transcription of the nisin gene cluster. This cluster includes genes involved in the production of premature Nisin-A and other peptides necessary for post-translational modification, nisin immunity, and signaling. As reported by Kuipers et al. (Kuipers et al., 1995), mutants of Nisin-A can still participate in the quorum sensing system, albeit to varying degrees. Given this background, it is plausible that NisinPV might interact with the quorum sensing mechanism in *Lactococcus lactis*. Such an interaction could potentially lead to an upregulation of gene expression within the nisin gene cluster, resulting in increased production of Nisin-A and related peptides. This interaction could be advantageous for the proposed therapeutic system, enhancing nisin production and potentially improving GMO’s antimicrobial properties. However, it’s essential to note that our study did not specifically investigate the interaction of NisinPV with the native quorum-sensing system of *Lactococcus lactis*. This aspect remains a potential avenue for future research to fully understand the implications of introducing NisinPV into this system.

A kill switch can be introduced that causes the GMO to lyse itself and prevents unintended gene transfer in the unlikely event that it does not come into contact with the particular pathogen. Using CRISPR Cas9 (Ran et al., 2013), the thyA gene in the chassis organism can be modified, making the mutant unviable in conditions with low levels of thymidine or thymine. A recessive deadly mutation called thyA renders the bacteria incapable of manufacturing any thymidine and forces them to rely on the environment for an external source of thymidine (Ross et al., 1990). The bacteria would die when thymidine is absent since they can no longer synthesize it. Thymidine can be added to the mutant bacteria while it is being cultivated in the bioreactor. After that, the vial can be kept at 4°C with thymidine added so that bacterial growth is minimal and the thymidine is not consumed. The bacteria could begin to proliferate after being introduced into the udder and using the thymidine. The lysis E7 lyses the chassis GMO and releases the bacteriocin in the event that it comes in contact with the target pathogen. The lack of thymidine in the udder’s environment will cause the chassis to perish even if it is not in contact with the pathogen. Hence, the GMO equipped with the kill switch would not be able to transfer its genes to any other bacteria residing in the microbiota of the udder and cause unprecedented problems and contamination.

Compared to traditional antibiotic treatments for cow mastitis, our method has a number of benefits. First off, it targets *Staphylococcus aureus* directly while expecting minimal damage to other microbes. The activity of NisinPV on the cattle’s udder microbiome needs to be experimentally validated(Derakhshani et al., 2018, Dean et al., 2021). *Staphylococcus aureus* is a key pathogen linked to bovine mastitis (Porcellato et al., 2020). Second, it prevents the over-use of antibiotics, thus reducing the development of antibiotic resistance. Third, the biofilm, which is a significant impediment to the penetration of antimicrobial drugs, may be more easily destroyed by DNaseI released by the GMO, than by antibiotics alone. Furthermore, because quorum sensing is a typical method utilized by many bacterial pathogens to coordinate pathogenicity and biofilm development, our strategy might be applicable to conditions other than bovine mastitis.

A probiotic strain called *Lactococcus lactis* LMG7930 is our choice of chassis organism to incorporate the proposed therapeutic system. The medication can be provided as a single dose, and it could be administered into the cow’s diseased udder via the teat canal via intramammary injection (Merriman et al., 2017). To thoroughly assess the safety interaction with native microbiome and effectiveness of this GMO-based treatment in vivo as well as the potential hazards connected with the discharge of genetically modified bacteria into the environment, additional research is required. However, this research offers a potential proof-of-concept for the creation of GMO-based antibiotics for bacterial infections, which could have broad ramifications for the antimicrobial therapy industry.

In our approach, we target *Staphylococcus aureus*, because it is responsible for almost half of the cases of Bovine mastitis (Birhanu et al., 2017). Mycobacteria as the primary causes of bovine mastitis are considered of low occurrence (Schiller et al., 2011). Commonly, it is the lipophilic species of the Corynebacterium genus that cause mastitis and they usually have prolonged incubation time and more meticulous or finicky growth requirements (Taylor et al., 2003).

The authors of this study acknowledge that all concentration measurements were conducted in *Escherichia coli*, and while our experiments specifically focused on the impact of DNaseI on biofilm formation in *Pseudomonas aeruginosa*, the study was not extended to or involved experiments with *Staphylococcus aureus* biofilm. It is well-documented that extracellular DNA (eDNA) is a significant component of biofilms formed by both *Pseudomonas aeruginosa* Thi et al., 2020 and *Staphylococcus aureus* Archer et al., 2011. Given that DNaseI is an enzyme known to degrade eDNA, our study is based on the working hypothesis of expecting similar outcomes with *Staphylococcus aureus* biofilm degradation as observed in our experiments with *Pseudomonas aeruginosa*. This working hypothesis is supported by the shared reliance of these bacterial species on eDNA as a structural component of their biofilms.

In conclusion, this paper presents a preliminary inquiry into a potential novel approach to utilize bioengineered defensins as a potent weapon against bovine mastitis. By unraveling the potential of this innovative strategy, we strive to contribute to the development of sustainable solutions for the dairy industry, ultimately safeguarding animal health, enhancing milk quality, and preserving human health.

## RESOURCE AVAILABILITY

All materials were sourced from organizations mentioned in the Funding section of the manuscript. Data generated from the work is available on the iGEM website. Data from Mathematical Modelling has been uploaded to GitHub.

## AUTHOR CONTRIBUTIONS

All the authors have contributed equally to this work.

## ACKNOWLEDGMENTS

The authors would like to thank Mr. Anshuman Jaysingh, Ms. Lakshmi Prakash, Ms. Soumi Bhattacharyya, Ms. Debshruti Biswas and Mr. Debdeep Chatterjee for their continuous support, guidance, and help during the work. Additionally, we would like to thank Prof. Supratim Dutta for his valuable insights and guidance. Finally, we would like to thank the Department of Biological Sciences, IISER Kolkata for helping us with lab equipment to carry out this project.

