## Supplementary Table 1 for "Exploring Quorum-Sensing-Based Bacterial GMOs for Addressing Bovine Mastitis: A Developmental Approach"

### 1. Key resources table :

| Reagent | Source | Additional information |
| --- | --- | --- |
| Bacterial strains : | | |
| ***E. coli* DH5α** | IISER Kolkata* | Used for cloning and assembling DNA |
| ***E. coli* BL21** |  | Used for studying protein expression |
| ***P. aeruginosa*** |  | Used for biofilm assay |
| Plasmids : | | |
| **pUC19** | NEB | Sponsored the project |
| **pSB1C3** | iGEM | Common vector for all the Biobricks present in iGEM-21 kit plates provided by iGEM headquarters. |
| Cell culture media : | | |
| LB ( Luria Broth ) premix | IISER Kolkata* |  |
| TSB (Tryptic Soy Broth) premix |  |  |
| NB ( Nutrient Broth ) premix |  |  |
| Reagents: | | |
| 0.5 M EDTA stock solution | IISER Kolkata* | Protocol for preparation of Stock solutions can be accessed from supplementary material. |
| 50X TAE Buffer |  |  |
| 10X TAE Buffer working solution |  |  |
| 1M NaOH solution |  |  |
| Crystal Violet solution |  |  |
| Antibiotics Stock and working solutions |  |  |
| TFB-I Buffer |  |  |
| TFB-II |  |  |
| DNAse I | Sigma - Aldrich | D2025-15KU ( Product identifier ) |
| EcoRI | Promega | R6011 ( catalog number ) |
| PstI | Promega | R6111( catalog number ) |
| Q5^®^ High-Fidelity 2X Master Mix | NEB | M0492S ( Catalog number, Used for normal PCR) |
| GoTaq® Green Master Mix | Promega | M7122 ( Catalog number, used for colony PCR) |
| T4 DNA Ligase | NEB | M0202M ( catalog number ) |
| T4 DNA Ligase Reaction Buffer | NEB | M0202M ( catalog number ) |
| Xbal | Promega | R6181 ( Catalog number ) |
| Spel | Promega | R6591 ( Catalog number ) |
| Commerical Kits/assays : | | |
| NEBuilder^®^ HiFi DNA Assembly Cloning Kit | NEB | E5520S( catalog number ) |
| Wizard® SV Gel and PCR clean up system kit | Promega | A9281 ( catalog number ) |
| Wizard® Plus SV Minipreps DNA purification System | Promega | A1330( catalog number ) |

*Provided by the Department of Biological Sciences, Indian Institute of Science Education and Research Kolkata.

- List of primers used is present below.
- List of DNA sequences ordered from Twist Bioscience is present below.
- Resource availability :

1. Materials availability : All materials were sourced from organisations mentioned in the Funding section of the manuscript.
2. Data and code availability : Data generated from the work is available in the iGEM website. Data from Mathematical Modelling has been uploaded into GitHub.

### 2. DNA Fragments ordered from Twist Bioscience :

**Eicsm6 Coding Region:**
GAATTCGCGGCCGCTTCTAGATGAAAATCCTGTTCTCTCCTATAGGAAATACGGACCCGTGGCGGAACGATCGAGACGGCGCAATGTTACATATTGTACGACATTATCAGCCCGATCGCGTTGTTTTGTTCTTCACTGAAAGCATATGGCAGGGCAACCAGCATTTCTCTGGGCAGCAGGCATTCGACTGGGTCAAGATTATACAATCAATCAATGAAAACTGCCAAATAGAGATCAAGTGCGACACGATAGAAGTCGAAAACGACTTTGACGCTTATAAGGACTTATTTCACCAATATCTCGTTGAAGAGAAGCGCAAGTACCCAAACGCCGAGATCTTTCTGAACGTAACCTCAGGTACCCCACAAATGGAGACGACACTGTGTCTCGAGTACGTAACCTATCCGGATAAGATGAGATGTATCCAAGTGAGTACTCCACTGAAAACCTCAAACGCGAAGACGAAGTATGCCCAGGCAGACTGTCAAGAGGTAGATTTGGAGATTGTAAACGAGGAAGAAAGCCAACAGCCGAGCCGGTGCCATAAAATTGCCATCCTCTCATTCAGAGAGGCCATTGTCAGAAACCAAATTAAGTCCCTGCTGGACAACTATGATTATGAGGCAGCTTTGCAGCTGGTCGCTAGTCAGAAGTCCTTTCGTAACGGTAAAGAGATCAGAAAGAAACTTAAGGAACTTATTGACGATATCAAAATGCACCGCGTTTTCAGTTATCTGATAAAGCAATACCCGCGTAATGAGAAACTTCAGAAGGCCCTTCTCCATACGATCCTGTTGGAGATGCGCCATCAGCGCGGCGATATAGCTGAGACATTAATTCGCGTTAAGTCAATCGCTGAGTACATTGTTGAGCAGTACATACAGAAGAATTACCCCTACCTCATTATTTACAAGGAAGATAAACCGTATTTTAATGTCTCGTATTCCCAGGAACTCACTGAATCGTACCTGGCGTTAATGGATTCCCGAAATAAGAAAACGAATAAGAAGATGACTGTAGATAGCCTCGACCGGATACTGGGATTCCCGGCCTATCGGGACTTTCTTCAGTTGCTCGAGGCGAGTAATGAAATGACAAATGAAATGAACAAGGTCAACGAGATCAACAATCTGCGGAACAAAGTGGCACACAATTTGGATAGCTTGAACCTCGACCGCGATAAGAACGGCCGTAAGATTACTAACGCAGTGACCGCTGTTCGTACTATGCTCCTTGCTGTGTTTCCTGAAGTGCAAGAGAATGATTTTCACTACCTCAAACAGTTCAACCAATCCATCAAGGAGCTGCTTTAATACTAGTAGCGGCCGCTGCAG

**Eicsm6 Composite part:**
GAATTCGCGGCCGCTTCTAGAGCAATACGCAAACCGCCTCTCCCCGCGCGTTGGCCGATTCATTAATGCAGCTGGCACGACAGGTTTCCCGACTGGAAAGCGGGCAGTGAGCGCAACGCAATTAATGTGAGTTAGCTCACTCATTAGGCACCCCAGGCTTTACACTTTATGCTTCCGGCTCGTATGTTGTGTGGAATTGTGAGCGGATAACAATTTCACACATACTAGAGAAAGAGGAGAAAATGAAAATCCTGTTCAGTCCTATCGGTAATACTGACCCCTGGCGTAACGATCGTGACGGAGCAATGCTTCATATTGTCCGCCATTATCAGCCCGATCGGGTTGTTCTGTTCTTCACAGAATCTATATGGCAGGGTAACCAGCATTTTAGCGGGCAGCAGGCATTCGACTGGGTTAAGATTATACAATCTATAAATGAAAATTGCCAAATAGAGATAAAGTGCGACACGATCGAAGTCGAAAACGACTTTGACGCGTATAAGGACCTCTTTCACCAATATCTTGTTGAAGAGAAGCGCAAGTACCCAAACGCCGAGATATTTCTTAACGTAACCAGTGGTACCCCACAAATGGAGACGACGCTGTGTTTAGAGTACGTCACCTATCCAGATAAGATGCGGTGTATACAAGTGTCAACGCCACTCAAAACCTCGAACGCAAAGACGAAGTATGCACAGGCTGACTGTCAAGAGGTCGATTTGGAGATTGTGAACGAGGAAGAAAGCCAACAGCCGTCCCGTTGCCATAAAATTGCCATACTCTCCTTCCGAGAGGCGATTGTGCGCAACCAAATTAAGAGCCTCTTGGACAACTATGATTATGAGGCTGCATTGCAATTAGTGGCCTCTCAGAAATCCTTTCGGAACGGAAAAGAAATACGTAAGAAATTGAAGGAACTTATAGATGATATCAAAATGCACCGCGTCTTTTCCTATCTGATTAAGCAATACCCTCGGAATGAAAAGTTGCAGAAGGCGTTACTTCATACTATCCTGCTGGAGATGCGCCATCAGCGGGGCGATATAGCGGAGACCTTGATTCGCGTTAAATCCATAGCCGAGTACATTGTAGAGCAGTACATTCAGAAGAATTACCCGTACCTTATAATTTATAAAGAGGATAAACCATACTTCAATGTGTCGTATAGCCAGGAATTAACCGAGTCATACTTAGCCCTCATGGATTCGCGTAATAAGAAAACGAATAAGAAGATGACGGTCGATTCATTAGACCGCATTCTGGGTTTCCCTGCCTATCGTGACTTTCTGCAACTCCTGGAAGCAAGTAATGAAATGACGAATGAAATGAACAAGGTGAACGAGATAAACAATCTGCGGAACAAAGTAGCACATAACTTAGACAGTCTTAATTTAGATCGTGACAAGAATGGTCGCAAAATCACGAATGCGGTAACTGCGGTGCGCACCATGCTGTTGGCGGTCTTCCCAGAGGTTCAGGAAAACGACTTCCATTATTTAAAGCAATTTAATCAGAGCATTAAAGAACTTTTGCATCACCATCACCATCACTAACCAGGCATCAAATAAAACGAAAGGCTCAGTCGAAAGACTGGGCCTTTCGTTTTATCTGTTGTTTGTCGGTGAACGCTCTCTACTAGAGTCACACTGGCTCACCTTCGGGTGGGCCTTTCTGCGTTTATATACTAGTAGCGGCCGCTGCAG

**DNaseI basic:**
GAATTCGCGGCCGCTTCTAGATGCGTGGAACAAGACTTATGGGATTACTCCTTGCGTTAGCAGGCCTTTTACAACTCGGGCTTAGTCTGAAAATTGCGGCATTTAATATTCGAACTTTCGGCGAAACGAAAATGTCAAACGCGACTTTAGCGTCCTATATAGTGCGAATAGTACGGCGCTATGATATTGTACTTATTCAAGAAGTACGTGATAGTCATTTGGTAGCCGTCGGCAAACTGCTTGATTACTTGAATCAAGACGATCCTAATACTTATCATTACGTTGTGAGCGAACCCTTAGGTCGTAATTCCTATAAGGAACGTTACCTTTTCTTGTTTCGTCCGAATAAAGTCTCGGTTTTGGATACGTATCAATATGATGATGGTTGTGAATCATGTGGTAATGATTCTTTTAGTCGCGAACCAGCCGTCGTGAAATTTTCTTCACATTCGACAAAAGTAAAAGAGTTCGCTATAGTGGCACTTCATTCTGCACCGTCCGATGCCGTCGCAGAAATCAACAGCCTGTATGACGTGTATCTCGACGTGCAACAAAAGTGGCATCTGAATGACGTTATGCTTATGGGAGACTTTAACGCCGATTGTTCCTATGTTACGTCAAGTCAATGGTCTAGTATACGTCTCCGGACCAGTAGCACATTTCAATGGTTAATCCCAGATTCCGCTGATACAACTGCGACCTCAACTAATTGTGCGTACGATCGCATTGTAGTTGCTGGAAGCCTTCTGCAATCTTCAGTAGTCCCAGGTTCAGCAGCCCCTTTCGACTTCCAAGCTGCATACGGACTGAGCAATGAGATGGCCCTGGCCATCAGTGACCATTACCCGGTGGAGGTGACGCTGACATAATACTAGTAGCGGCCGCTGCAG

**DNaseI composite:**
GAATTCGCGGCCGCTTCTAGAGCAATACGCAAACCGCCTCTCCCCGCGCGTTGGCCGATTCATTAATGCAGCTGGCACGACAGGTTTCCCGACTGGAAAGCGGGCAGTGAGCGCAACGCAATTAATGTGAGTTAGCTCACTCATTAGGCACCCCAGGCTTTACACTTTATGCTTCCGGCTCGTATGTTGTGTGGAATTGTGAGCGGATAACAATTTCACACATACTAGAGAAAGAGGAGAAAATGCGGGGTACACGTTTGATGGGTTTACTCCTGGCTTTGGCCGGCTTGCTCCAATTGGGACTTAGTCTTAAAATTGCCGCGTTTAATATACGTACATTCGGCGAAACTAAAATGTCTAACGCCACATTGGCTTCTTATATCGTGCGTATTGTTAGACGATATGATATTGTGTTGATTCAAGAAGTTCGTGATAGTCATTTGGTTGCCGTCGGCAAACTTTTAGATTACCTTAATCAAGACGATCCGAATACATATCATTACGTTGTGTCCGAACCTTTGGGGAGAAATTCTTATAAAGAACGGTATCTTTTCCTGTTTCGACCGAATAAAGTTAGCGTACTCGATACTTATCAATATGATGATGGGTGTGAAAGCTGTGGCAATGATTCTTTTAGCCGTGAACCGGCAGTCGTAAAATTTAGTTCACATTCGACTAAAGTGAAAGAGTTCGCAATAGTCGCATTACATTCTGCACCGTCTGATGCTGTCGCGGAAATCAACTCGTTATATGACGTGTATTTAGACGTTCAACAGAAATGGCATTTAAATGACGTTATGCTGATGGGTGACTTTAACGCGGATTGTTCTTATGTCACGTCTTCTCAATGGTCTAGTATTAGACTTCGGACTTCAAGTACTTTTCAATGGTTAATCCCAGATAGCGCAGATACAACCGCCACCTCAACCAATTGTGCTTACGATCGGATAGTCGTTGCGGGTAGTCTCTTACAATCTAGCGTTGTCCCGGGATCAGCGGCCCCGTTCGATTTTCAGGCCGCTTATGGTTTGAGTAACGAAATGGCACTTGCAATTAGCGATCACTATCCCGTTGAAGTAACCTTGACCTAACCAGGCATCAAATAAAACGAAAGGCTCAGTCGAAAGACTGGGCCTTTCGTTTTATCTGTTGTTTGTCGGTGAACGCTCTCTACTAGAGTCACACTGGCTCACCTTCGGGTGGGCCTTTCTGCGTTTATATACTAGTAGCGGCCGCTGCAG

**NisinPV Coding Sequence:**
GTGCAACCTTGGGTAGGCTTTACTTAGCTAAAGGCCTTAGGGGATGGAATTCGCGGCCGCTTCTAGATGTCGACCAAGGACTTCAATCTGGACCTTGTTAGCGTCAGTAAGAAGGACTCCGGAGCGTCTCCGCGGATAACGTCAATATCTCTTTGCACCCCTGGCTGCAAGACTGGCGCGCTCATGGGATGCAATATGAAGACTGCGACCTGCCACTGCCCGGTCCATGTCTCTAAGTAATACTAGTAGCGGCCGCTGCAGCATCGTTTACTTGACTGAAAAGGGGCCTTCTCAAATTGG

**P2 + GFP :**
ATTAAATACAAATTACATTTAACAGTTAAGTATTTATTTCCTACAGTTAGGCAATATAATGATAAAAGATTGTACTAAATCGTATAATGACAGTGAAAAGAGGAGAAAATGCGCAAGGGCGAGGAGTTGTTTACAGGCGTGGTTCCGATCTTGGTCGAGCTTGACGGAGACGTAAACGGTCATAAGTTCAGCGTGTCCGGGGAAGGCGAGGGCGACGCTACCTATGGCAAGTTAACGCTGAAGTTCATCTGTACGACCGGTAAGCTTCCGGTCCCCTGGCCTACTTTGGTTACAACCTTTGGATACGGGGTGCAGTGTTTCGCACGATATCCGGACCACATGAAGCAACACGATTTCTTTAAATCAGCAATGCCAGAGGGATACGTTCAAGAGCGCACCATTTTCTTTAAGGACGATGGCAATTATAAAACGAGAGCAGAGGTAAAATTCGAGGGAGACACGTTAGTGAACCGAATTGAATTGAAGGGAATCGACTTCAAAGAGGACGGCAATATCCTCGGCCATAAGTTAGAGTATAATTACAATAGTCATAACGTGTACATAATGGCTGATAAGCAGAAGAACGGCATTAAGGTAAACTTTAAGATCCGTCATAACATCGAGGACGGCTCGGTACAGCTGGCTGATCACTACCAGCAGAACACGCCTATCGGAGACGGACCGGTTTTGCTTCCTGATAATCACTATCTTTCTACCCAGAGTGCATTGTCAAAGGACCCTAATGAGAAACGCGATCATATGGTATTGTTGGAGTTCGTCACCGCAGCGGGCATAACTCACGGTATGGACGAGCTGTATAAGTAATAACCAGGCATCAAATAAAACGAAAGGCTCAGTCGAAAGACTGGGCCTTTCGTTTTATCTGTTGTTTGTCGGTGAACGCTCTCTACTAGAGTCACACTGGCTCACCTTCGGGTGGGCCTTTCTGCGTTTATAGCTTTCGAAGGCTTAGGCGGGAG

**AgrA + terminator:**
AAAGAGGAGAAATACTAGATGGAGATTGCCTTGGCGACTGACAACCCCTACGAAGTATTAGAGCAAGCGAAGAATATGAATGATATTGGTTGTTACTTCCTTGACATCCAATTATCCACAGATATAAATGGAATAAAGCTCGGAAGCGAGATACGCAAGCACGACCCAGTGGGCAATATCATCTTTGTCACGTCCCACTCCGAATTGACATACTTAACCTTTGTGTATAAAGTGGCCGCGATGGACTTTATATTCAAGGATGACCCTGCCGAATTGCGCACACGTATAATTGATTGCTTAGAGACAGCTCATACTCGGCTTCAGTTGCTTAGCAAGGACAATAGCGTTGAGACAATCGAGCTTAAGCGCGGGAGTAATTCGGTGTATGTGCAGTACGACGACATTATGTTCTTTGAGAGTTCTACAAAATCTCATCGCTTAATTGCACACCTTGACAACAGACAAATAGAATTTTACGGAAACCTTAAGGAATTATCACAACTGGACGACAGATTCTTTCGGTGCCACAATAGTTTTGTGGTGAACAGACATAATATTGAGAGCATTGATTCAAAGGAAAGAATAGTCTATTTTAAGAATAAAGAGCATTGCTACGCAAGCGTGCGGAACGTAAAGAAGATTTAATAATACTAGAGCCAGGCATCAAATAAAACGAAAGGCTCAGTCGAAAGACTGGGCCTTTCGTTTTATCTGTTGTTTGTCGGTGAACGCTCTCTACTAGAGTCACACTGGCTCACCTTCGGGTGGGCCTTTCTGCGTTTAT

**pBAD + AgrC:**
ACATTGATTATTTGCACGGCGTCACACTTTGCTATGCCATAGCAAGATAGTCCATAAGATTAGCGGATCCTACCTGACGCTTTTTATCGCAACTCTCTACTGTTTCTCCATACCGTTTTTTTGGGCTAGCAAAGAGGAGAAAATGATCCTTATGTTTACAATCCCTGCGATCATCTCAGGAATCAAATATTCAAAATTGGACTACTTCTTTATTATTGTTATAAGTACGTTATCATTATTCCTCTTTAAGATGTTTGACAGTGCTTCTTTGATAATTCTCACCAGCTTTATTATTATTATGTACTTTGTGAAGATCAAGTGGTACTCGATTCTCCTGATAATGACAAGCCAAATAATTTTGTATTGCGCAAATTATATGTACATAGTAATTTATGCATATATTACTAAGATTTCTGACTCAATATTTGTCATTTTCCCAAGCTTCTTCGTAGTATACGTAACTATATCGATCTTATTCAGTTACATCATTAACCGGGTATTGAAGAAAATTTCCACACCCTATTTAATCTTAAATAAGGGGTTCTTGATTGTTATTTCAACAATATTGCTCTTGACGTTTAGCCTTTTCTTCTTCTATTCCCAAATAAATTCGGACGAAGCGAAGGTGATTAGGCAATATTCTTTTATTTTCATCGGCATAACAATTTTCCTCTCAATCCTCACGTTTGTGATTAGCCAATTTCTTCTGAAGGAGATGAAATATAAACGGAATCAGGAAGAGATTGAAACATACTACGAATATACGTTGAAAATAGAGGCTATTAATAATGAGATGCGGAAGTTCCGCCATGATTATGTCAATATCCTTACGACGCTTTCAGAGTATATTCGCGAGGATGATATGCCGGGATTACGCGACTACTTTAATAAGAATATAGTACCTATGAAGGACAATCTTCAGATGAATGCCATCAAATTAAATGGCATTGAGAATCTTAAGGTTCGGGAAATTAAAGGCTTGATAACAGCAAAGATTTTGAGAGCCCAAGAGATGAATATTCCGATCTCAATTGAGATTCCGGATGAGGTGAGCTCGATCAATTTAAATATGATAGATTTAAGCAGATCAATTGGGATTATTTTGGATAATGCAATTGAGGCGAGCACAGAGATTGATGATCCTATCATCCGTGTAGCGTTCATTGAATCGGAAAATTCGGTCACATTCATTGTCATGAATAAGTGTGCCGATGACATTCCTCGAATCCATGAGCTCTTTCAAGAGTCCTTTTCCACAAAAGGCGAGGGCAGAGGGCTCGGGTTGAGTACATTGAAGGAAATTGCAGACAATGCTGATAATGTGCTCTTGGATACAATAATTGAGAATGGCTTCTTCATCCAAAAGGTTGAGATTATTAACAATTAATAA

### 3. Primers :

**Primers for PCR:**

- **Amplification of biobricks :**

| **Primer Type** | **Sequence** | **%GC** | **TM** | **Length** |
| --- | --- | --- | --- | --- |
| **Forward** | **5’-gatggaattcgcggccgcttcta-3’** | **56.5** | **61.7** | **23** |
| **Reverse** | **5’-gatgctgcagcggccgctactagta-3’** | **60** | **64.4** | **25** |

- **Linearized plasmid backbone synthesis:**

| **Primer Type** | **Sequence** | **%GC** | **TM** | **Length** |
| --- | --- | --- | --- | --- |
| **Forward** | **5’- gtgctgcagtccggcaaaaaa-3’** | **52.4** | **59.3** | **21** |
| **Reverse** | **5’-gtgaattccagaaatcatccttagcg-3’** | **42** | **66** | **26** |

- **Sequencing primers :**

| **Primer Type** | **Sequence** | **%GC** | **TM** | **Length** |
| --- | --- | --- | --- | --- |
| **Forward (VF2)** | **5’-tgccacctgacgtctaagaa-3’** | **50** | **60** | **20** |
| **Reverse (VR)** | **5’-attaccgcctttgagtgagc-3’** | **50** | **60** | **20** |

**Primers specific for Hi-Fi DNA assembly:**

- For Unidirectional Assembly

| Pbad_RBS_AgrC_fwd_uni | 5’-tctggaattccaacattgattatttgcacgg-3’ |
| --- | --- |
| Pbad_RBS_AgrC_rev_uni | 5’-tttctcctctttttattaattgttaataatctcaaccttttg-3’ |
| RBS_AgrA_fwd_uni | 5’ taacaattaataaaaagaggagaaatactagatg 3 |
| RBS_AgrA_rev _uni | 5’ tttgtatttaattataaacgcagaaaggcc 3 |
| P2_GFP_fwd_uni | 5’ttctgcgtttataattaaatacaaattacatttaacagttaagtatttatttcctacagttag 3’ |
| P2_GFP_rev_uni | 5’ ggactgcagcacctcccgcctaagccttcg 3’ |
| PSB1C3_fwd_uni | 5’ gcttaggcgggaggtgctgcagtccggcaaaaaag 3’ |
| PSB1C3_rev_uni | 5’ aaataatcaatgttggaattccagaaatcatccttagcg 3’ |

- For Bidirectional Assembly

| Pbad_RBS_AgrC_fwd_bi | 5’- tttgtatttaatacattgattatttgcacgg-3’ |
| --- | --- |
| Pbad_RBS_AgrC_rev_bi | 5’-tttctcctctttttattaattgttaataatctcaaccttttg-3’ |
| RBS_AgrA_fwd_bi | 5’ taacaattaataaaaagaggagaaatactagatg 3 |
| RBS_AgrA_rev _bi | 5’ ggactgcagcactataaacgcagaaaggcc 3 |
| P2_GFP_fwd_bi | 5tctggaattccactcccgcctaagccttcg 3’ |
| P2_GFP_rev_bi | 5’aaataatcaatgtattaaatacaaattacatttaacagttaagtatttatttcctacagttag 3’ |
| PSB1C3_fwd_bi | 5’ ttctgcgtttatagtgctgcagtccggcaaaaaag3’ |
| PSB1C3_rev_uni | 5’ gcttaggcgggagtggaattccagaaatcatccttagcg 3’ |

### 4. Stock Solution preparation :

**0.5 M EDTA stock solution:**

For preparation of 30ml stock solution :

- Weigh out 5.583 grams of EDTA disodium salt.
- Dissolve in 25ml of milli Q water and adjust pH with solid NaOH to reach the pH of 8.
- Add 5ml milli Q water to make the total volume to 30ml.
- Autoclave for 15mins at 121 °C.

**50X TAE Buffer:**

For preparation of 250ml solution:

- Weigh out 60.5g Tris base and dissolve approximately in 200ml milli Q water.
- Carefully add 14.275ml of 100% glacial acetic acid.
- Add 25ml of 0.5 M EDTA (pH 8).
- The pH of buffer should be around 8.5.
- Store at room temperature.

**10X TAE Buffer working solution:**

Add 20ml of 50X TAE buffer to 1000ml of milliQ water.

**1M NaOH preparation for pH adjustments:**

To prepare 20ml of 1M NaOH add 0.8g of NaOH pellets in 20ml of dH2O. Then store it in a falcon tube.

**Crystal Violet solution preparation:**

For 1L solution:

1. Add 1g of crystal violet to 1000ml of dH2O and then do the magnetic stirring for solubilising.

**Antibiotics Stock and working solutions:**

As per the requirement, the stock and working concentrations can be made by referring to the corresponding concentrations provided in the table below.

| ANTIBIOTIC | CONCENTRATION OF STOCK SOLUTION | STORAGE TEMP. | WORKING CONCENTRATION |
| --- | --- | --- | --- |
| Ampicillin | 100 mg/ml (H2O) | –20°C | 100 μg/ml |
| Carbenicillin | 100 mg/ml (H2O) | –20°C | 20–200 μg/ml |
| Chloramphenicol | 34 mg/ml (ethanol) | –20°C | 30 μg/ml |
| Kanamycin | 10 mg/ml (H2O) | –20°C | 10–50 μg/ml |
| Streptomycin | 10 mg/ml (H2O) | –20°C | 10–50 μg/ml |
| Tetracycline | 5 mg/ml (ethanol) | –20°C | 10–50 μg/ml |

**Stock for Primers:**

- Check the concentrations of each primer provided by the vendor.
- Need to make the stock concentration of 100pm/μl.
  - Centrifuge the empty vials at 30,000 rpm for 30 secs.
  - Heat the NFW at 95°C for 10mins then at 55°C for 30 mins
  - Pour the NFW to the mentioned volume to make the stock concentration (100pm/μl) in each tube of primer. Remember to perform this step inside the hood.
- Leave the tubes in 4°C for 4-5 hrs.
- Take it out from 4°C and mix by soft pipetting or light vortex. Then store it at -20°C.
- For making a working concentration of 10pm/μl add 10μl of stock solution to 90 μl of NFW.

**Resuspension of biobricks from iGEM kit Plates**

- Label the plates properly and carefully with a marker (for reference check the well contents from iGEM website)
- Take 10μl of NFW using a micropipette. Punch a hole in the aluminum foil for the well you want to resuspend with the tip of the micropipette tip.
- Slowly release the water into the well.
- Gently pipette inside the well for a few times. The solution will be red due to the presence of cresol dye.
- Keep the mixture there for 5 mins
- Now transfer it to a PCR tube and store it in -20°C. Label it properly before storing

**Resuspension of DNA fragments ordered from Twist Bioscience -**

- Briefly centrifuge the tube and resuspend the DNA in NF-tris EDTA (TE Buffer), pH 8 or 10mM Tris HCl, pH 8 to the desired concentration.
- A concentration of 10ng/μl is recommended for a stock solution.
- Prepare aliquots of the stock and make working aliquots.
- For long-term storage, freeze DNA at -20°C or -80°C

**DNaseI stock preparation**

Lyophilized form of Bovine DNaseI was obtained from sigma aldrich (D2025-15KU). Resuspension and stock preparation was done as per the guidelines from the company:

1. Lyophilized form which contains approximately 5 mg of DNaseI was resuspended in 0.15M NaCl solution making up to a concentration of 5mg/ml.
2. Working concentrations were made by diluting it further in 0.15M NaCl or Media directly.
3. Stock solution was stored in -20 ℃ to prevent degradation.

**TFB-1(pH-5.1) composition for 30 ml :**

- 10 mM RbCl - 0.36 g
- 50 mM MnCl2 - 0.19 g
- 30 mM CH3COOH+ - 0.08 g
- 100 mM CaCl2 - 0.03 g
- 15 % glycerol - 4.5 ml

**TFB-2(pH-6.8) composition for 15 ml**

- 10 mM MOPS -0.031 g
- 10 mM RbCl-0.012 g
- 15 mM CaCl2-0.124 g
- 15 % glycerol-2.25 ml

### 5. Figures

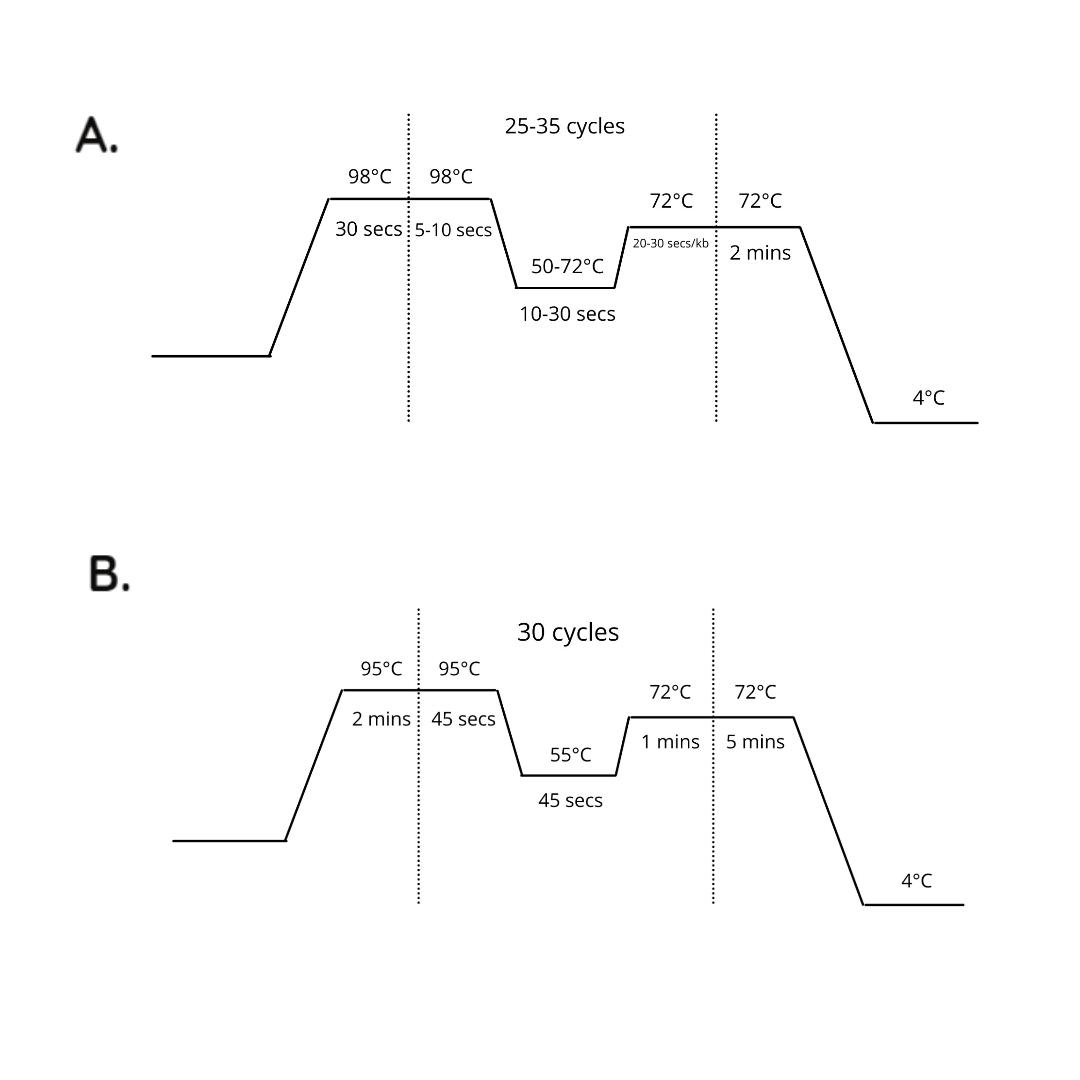

Figure 1 : A. Normal PCR , B. Colony PCR.

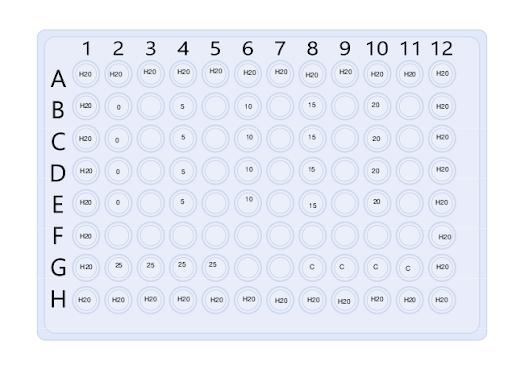

Figure 2 : Design of 96 Well Plate
